# Molecular response to the non-lytic peptide bac7 (1-35) triggers disruption of *Klebsiella pneumoniae* biofilm

**DOI:** 10.1101/2025.08.08.669280

**Authors:** Robert L. Beckman, Berta Martinez, Flor Z. Santiago, Gabriela N. Echeverria, Bruno V. Pinheiro, Marcelo D.T. Torres, Logan Suits, Shantal Garcia, Paeton L. Wantuch, Cesar de la Fuente-Nunez, Prahathees Eswara, David A. Rosen, Renee M. Fleeman

## Abstract

*Klebsiella pneumoniae* is becoming increasingly difficult to treat as multidrug-resistant (MDR) strains become more prevalent. The formation of biofilm heightens this threat by embedding bacterial cells in a polysaccharide-rich matrix that limits antibiotic penetration. Here we dissect the anti-biofilm bovine host-defense cathelicidin peptide fragment bac7 (1-35), exploring its anti-biofilm mechanism, evaluating its ability to curb dissemination of hypervirulent *K. pneumoniae*, and testing its breadth of activity against diverse clinical isolates. Transcriptomic profiling revealed that bac7(1-35) simultaneously compromises the bacterial membrane and inhibits ribosomal function, a dual assault that precipitates rapid biofilm collapse and blocks bacterial spread. Further, the peptide eradicated biofilms produced by the strongest MDR clinical isolates in the Multidrug-Resistant Organism Repository and Surveillance Network (MRSN) diversity panel. Although bac7 (1-35) kills bacterial cells via a cytosolic mechanism, membrane interaction profiles varied among MRSN isolates, correlating with differential peptide translocation. In a delayed-treatment murine skin-abscess model, bac7 (1-35) halted *in vivo* dissemination of the hypervirulent strain NTUH-K2044. Collectively, these results delineate a multifaceted mode of action for bac7 (1-35) and underscore its therapeutic promise against biofilm-associated MDR *K. pneumoniae* infections.

**Author Summary:** *Klebsiella pneumoniae* is a top-priority pathogen for new therapies, with many strains already approaching pan-drug resistant status. Biofilm formation further complicates treatment, yet biofilm-active therapeutics have not reached the clinic, in part because we still lack a detailed understanding of how to disrupt these impenetrable structures. Antimicrobial peptides are promising candidates and have shown biofilm-disruption potential. Here we provide mechanistic insight into how a host defense peptide dismantles pre-formed *K. pneumoniae* biofilms. We find that the peptide’s dual targeting of bacterial membranes and ribosomes triggers dispersal from the biofilm state and concomitantly downregulates factors required for surface attachment and extracellular matrix production. This mechanism involves a protein that, to our knowledge, has not been characterized in *K. pneumoniae*. Our findings reveal a switch that can be leveraged to reprogram biofilm maintenance toward dispersal in *K. pneumoniae*, advancing the path to peptide-based antibiofilm therapeutics.

## Introduction

*Klebsiella pneumoniae* has emerged as a priority pathogen in recent years due to its extreme drug resistance and the emergence of virulence traits ^1–8^. Over the past few decades, two distinct pathotypes have been reported: classical *K. pneumoniae* (cKp), many of which are multi-drug-resistant (MDR) and commonly infect patients with underlying chronic diseases; and hypervirulent *K. pneumoniae* (hvKp), which can afflict healthy individuals in the community ^5–7^. Regardless of pathotype, biofilm formation is associated with 60-80% of hospital-associated infections and increases the antibiotic resistance and immune evasion of isolates that may otherwise be susceptible ^9^. This complex matrix is often impenetrable to standard antibiotics and protects the embedded bacterial cells from the immune system, resulting in recalcitrant hospital-associated infections ^9^. Despite the significance of these infections, effective strategies to penetrate and disrupt biofilms remain an unmet need.

Biofilm formation is essential for bacterial survival under environmental stress and is a highly regulated process involving transcriptional changes guided by environmental stressors and signal responses ^10, 11^. During infection, environmental stimuli, such as limited nutrients, temperature, and change in pH, promote biofilm formation and persistent infections ^11, 12^. Signaling molecules that respond to environmental stressors act as molecular switches to coordinate the transcriptional changes necessary for biofilm formation ^12, 13^. Fimbriae and exopolysaccharides have been shown to be important for biofilm formation and regulated by the second messenger cyclic diguanosine monophosphate (c-di-GMP) ^14–16^. Due to its important role in biofilm formation, c-di-GMP has been viewed as a promising drug target for combating biofilm formation, which can be targeted directly or indirectly by interfering with cellular processes involved in its regulation ^12^. However, targeting c-di-GMP directly may prove challenging as there are multiple diguanylate cyclase enzymes with redundant abilities to produce c-di-GMP ^17^.

Antimicrobial peptides have shown promise as therapeutics targeting MDR Gram-negative pathogens and have displayed the potential to disrupt biofilms ^18–20^. Mammalian neutrophil derived non-lytic polyproline host-defense peptides are unique, as they exert intracellular antimicrobial effects without causing membrane lysis like most antimicrobial peptides ^21, 22^. Instead, their primary mode of action is the inhibition of protein translation through binding the 50S ribosomal subunit ^23–25^. Entry into the cytoplasm is typically mediated by stereospecific binding to the SbmA transporter ^25^. However, the mechanism of action of bac7 (1-35), henceforth referred to as bac7 varies between bacterial genera, with membrane-targeting activity observed in *Pseudomonas aeruginosa* and intracellular inhibition documented in *Salmonella enterica* and *Escherichia coli* ^24^. Our previous work has shown that bac7 binds polysaccharides and induces biofilm collapse in hvKp NTUH-K2044 ^26^. Importantly, the decreased mucoidy in treated biofilm supernatant highlights the potential of bac7 to decrease the hypermucoviscous nature of hvKp and suggests it might prevent *in vivo* dissemination of hvKp. However, the contribution of bac7’s membrane interactions to its mechanism of action against *K. pneumoniae* remains unclear.

Here, we demonstrate that bac7 induces both membrane stress and ribosome inhibition, embodying a dual mechanism of action against *K. pneumoniae*. Transcriptional analysis revealed a membrane stress response without cell lysis, and activation of *mgtC*-mediated biofilm release by modulating c-di-GMP activated cellulose and type I fimbriae. Expanding our analysis using the Multidrug-Resistant Organism Repository and Surveillance Network (MRSN) *K. pneumoniae* diversity panel ^27^, we found variable bac7 induced membrane depolarization amongst diverse isolates and heterogeneous peptide uptake within single populations. Testing bac7 using an *in vivo* skin abscess model revealed bac7 treatment has the potential to decrease dissemination of hvKp NTUH-K2044. Our results show that the dual mechanism of action of bac7 towards *K. pneumoniae* triggers a molecular response to decrease cellulose and fimbriae leading to biofilm disruption and decreased dissemination.

## Results

### Transcriptional insight reveals complex mechanism of killing and biofilm release

There has been a diversity of reported mechanisms of bacterial killing by bac7, and these mechanisms have been shown to vary between bacterial genera ^24, 28^. Given the multiple potential mechanisms, we aimed to identify the mechanism of killing in hvKp and to understand the rapid biofilm collapse that occurs before viability changes observed in our original study ^26^. We hypothesized that there is a molecular mechanism of peptide-mediated biofilm release. We therefore investigated the molecular response of hvKp to bac7 using RNA sequencing of hvKp NTUH-K2044 with 7.5 µmol L^−1^ of bac7 for 30 minutes.

We found a total of 2,511 genes that displayed significant differential expression (-log10(FDR) > 2) following treatment with bac7 compared to the no treatment control group, with 1,202 genes upregulated (log2 Fold Change > 1) and 1,309 genes downregulated (log2 Fold Change < −1) (**Supplemental data file 1**). To provide context to the overall gene expression changes with bac7, we used KEGG and STRING database analyses ^29–31^. The KEGG pathway enrichment analysis revealed ribosome (54 genes upregulated, p-value 1.5e^−21^), oxidative phosphorylation (37 genes downregulated, p-value 7.5e^−10^), and the TCA cycle (28 genes downregulated, p-value 1.6e^−8^) pathways were the most changed with treatment (**Supplemental Figure 1**; **Supplemental Table 1**). When considering gene ontology, we found genes associated with energy production were again the most enriched (**Supplemental Figure 2**). These pathways have been shown to be common metabolic responses to antibiotic treatment but also antimicrobial peptides ^32, 33^. The STRING protein-protein interaction network analysis (FDR <1 e^−10^, LogFC >1.5) revealed significantly more interactions than expected indicating a potential biological connection between the differentially expressed genes (**Supplemental data file 2**). We found two large clusters of interacting proteins corresponding to protein translation (increased expression) and oxidative phosphorylation (decreased expression), with F_O_F_1_ ATP synthase subunit epsilon (KP1_RS25675) serving as the connecting node between these groups (**Supplemental Figure 3**). Interestingly, we found the ribosomal cluster connected with transcription factors to reveal a footprint of membrane stress. Specifically, *rpoB* and *rpoC* genes encoding for RNA polymerase subunits were decreased in expression with bac7 (−2.5 LogFC, p-value 1.9e^−12^ and −4.1 LogFC, p-value 1.5e^−14^, respectively) indicating potential overall decrease in transcription levels, while *rpoE* the gene encoding for the membrane stress response transcription factor was increased in expression (+3.2 LogFC, p-value 5.2e^−13^), suggesting membrane stress is induced by bac7. Looking downstream at RpoE regulated genes, we found *ompC* to have decreased expression (−4.0 LogFC, p-value 6.0e^−15^), possibly in response to the increased expression of *rseA* (+3.0 LogFC, p-value R 2.4e^−13^), encoding for an anti-sigma factor that blocks RpoE activity. Our data suggest that the bac7 mechanism of killing *K. pneumoniae* involves both ribosome inhibition and membrane stress.

When looking at the individual genes displaying the most significant changes in expression (-log10 FDR > 1.0e^−10^) following treatment with bac7, we found changes not only indicating membrane and ribosomal stress but also changes suggesting a switch out of biofilm formation state (**Figure 1A**). Specifically, polyamines have been shown to be important for biofilm formation ^34^, and lysine decarboxylase (*cadA*), important for polyamine cadaverine production displayed the greatest decrease in expression of all genes in our profile (−9.3 LogFC, p-value 5.5e^−16^) ^35^. We also saw decreased expression of the genes involved in export of the polyamine putrescine (*sapCDF* −5.7 LogFC, p-value 1.6e^−13^; −4.5 LogFC, p-value 6.4e^−13^; and −6.3 LogFC, p-value 1.5e^−13^, respectively). In addition, we found increased expression of toxin-antitoxin system genes (*vapBC +*4.6 LogFC, p-value 2.5e^−12^; +5.1 LogFC, p-value 2.4e^−07^), shown to be involved in resistance to antimicrobial peptides and biofilm formation ^32^. Finally, we found significant increased expression of *mgtC* (MgtC family protein +5.4 LogFC, p-value 3.6e^−16^), that encodes a protein shown to inhibit FoF1 ATP synthase and decrease c-di-GMP availability ^36^. Intriguingly, we found bac7 treatment significantly decreased expression of FoF1 ATP synthase genes (*atpACDG* −4.6 LogFC, p-value 3.0e^−14^; −5.6 LogFC, p-value 2.3e^−15^; −5.9 LogFC, p-value 1.2e^−15^; and −5.4 LogFC, p-value 7.9e^−15^, respectively). Furthermore, the most significantly upregulated gene in our analysis (KP1_RS24555 +6.8 LogFC, p-value 4.4e^−17^) encodes for an ATP-binding cassette domain-containing protein that when blasted on NCBI shows 99% protein homology to EttA (GenBank: CAH5943538.1), that acts as an energy-dependent translational throttle shown to respond to cellular changes in ATP ^37^. When looking more broadly beyond the stringent false discovery rate cutoff, we found the c-di-GMP activated type 1 fimbriae and cellulose operons ^16, 38^ (*fimACDGHK*, *bcsA1B1CFGZ, and bcsA2B2*) had decreased expression with bac7 treatment (**Table 1**).

**Figure 1.**
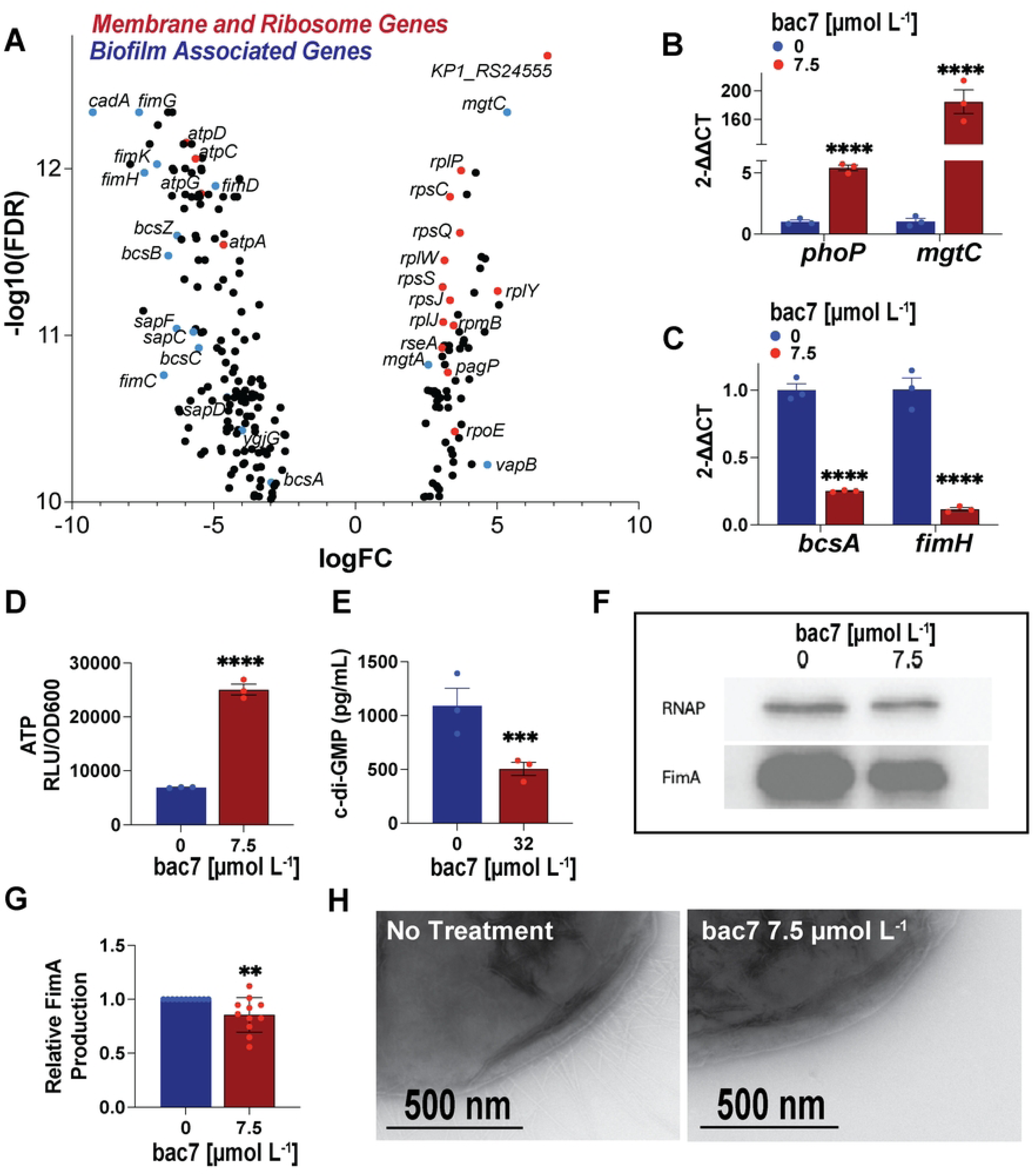
Transcriptional insight reveals complex mechanism of killing and biofilm release. Volcano plot of *K. pneumoniae* NTUH-K2044 most significantly differential expressed genes (DEGs) (-log_10_ FDR >10) following 7.5 µmol L^−1^ bac7 for 30 minutes (**Figure 1A**). Red dots show gene changes relevant to the mechanism of killing and blue dots show biofilm important gene changes. RT-qPCR validation of *phoP* and *mgtC* (**Figure 1B**), and c-di-GMP regulated *bcsA* and *fimH* (**Figure 1C**) with 7.5 µmol L^−1^ bac7 for 30 minutes. Intracellular ATP assessment following 7.5 µmol L^−1^ bac7 for 30 minutes measuring relative luminescence units (RLU) using a CellTiter-Glo^®^ kit and normalized to the respective OD_600_ (RLU/OD_600_) (**Figure 1D**). Intracellular c-di-GMP levels using a cyclic di-GMP ELISA kit (**Figure 1E**). The RT-qPCR was normalized using a *ftsZ* housekeeping gene. One-way ANOVA was used to determine significance compared to the no treatment groups for **Figures 1B-1E** with Dunnett’s correction for multiple comparisons with adjusted p-values shown (asterisks indicate p-values **** <0.0001, ** <0.01, and ns >0.1) and error shown reported as ± SEM. Immunoblot of hvKp NTUH-K2044 with 7.5 µmol L^−1^ bac7 treatment showing FimA antibody staining next to RNA polymerase alpha subunit control (**Figure 1F**). The relative abundance of FimA from n=11 western blots with error shown as ± SD and significance determined by unpaired t-test (p-value 0.0076) (**Figure 1G**). Transmission electron microscopy of hvKp NTUH-K2044 with or without 7.5 µmol L^−1^ bac7 using antibody labeling of FimA fimbriae (**Figure 1H**).

**Table 1.** Bcs and fim operons decreased expression with bac7 treatment.

| Gene | Description | Log2<br>FC | pValue |
| --- | --- | --- | --- |
| *fimG* | Tip adaptor subunit FimG | -7.6 | 4.9E-14 |
| *fimH* | Adhesion protein FimH | -7.4 | 5.1E-15 |
| *fimK* | EAL domain-containing protein FimK | -7.0 | 2.8E-15 |
| *fimC* | Periplasmic chaperone protein FimC | -6.7 | 4.0E-13 |
| *fimD* | Fimbrial biogenesis usher protein FimD | -4.9 | 6.8E-15 |
| *fimA* | Type 1 fimbrial major subunit FimA | -1.3 | 2.5E-09 |
| *bcsB* | Cyclic di-GMP-binding regulatory protein BcsB | -6.6 | 3.6E-14 |
| *bcsZ* | Periplasmic endoglucanase BcsZ | -6.3 | 2.5E-14 |
| *bcsC* | Outer membrane protein BcsB | -5.5 | 2.4E-13 |
| *bcsG* | Cellulose biosynthesis protein BcsG | -3.9 | 7.7E-12 |
| *bcsA* | UPD-forming cellulose synthase catalytic subunit | -3.2 | 5.6E-12 |
| *bcsB2* | Cyclic di-GMP-binding regulatory protein BcsB | -3.0 | 4.5E-11 |
| *bcsA2* | UPD-forming cellulose synthase catalytic subunit | -2.9 | 3.3E-12 |
| *bcsF* | Cellulose biosynthesis protein BcsF | -2.6 | 6.5E-07 |

We hypothesized the decreased expression of cellulose and fimbriae operon genes are due to the increased expression of the gene encoding for the MgtC family protein. There are no studies of this protein in *K. pneumoniae*; however in *S. enterica,* expression of *mgtC* is controlled by the global response regulator PhoP (*phoP* 1.3 LogFC, p-value 1.6e^−08^) in response to membrane stress ^38, 39^. Once transcribed, *mgtC* upstream hairpin loops allow for autoregulation in response to elevated ATP under stressed conditions. When cells are treated with a ribosomal inhibitor, MgtC inhibits F_O_F_1_ ATP synthase to protect the cell from ATP toxicity caused by inactive ribosomes ^40^, consequently depleting c-di-GMP, which is an allosteric activator of cellulose and fimbriae expression (type 1 and 3) ^16, 38^. We used RT-qPCR to validate the bac7 expression changes in *phoP* and *mgtC*, as well as the downstream *bcsA* and *fimH*, known to be transcriptionally activated by c-di-GMP ^38^. Repeating the bac7 treatment used for RNA sequencing revealed significant increased expression of both *phoP* and *mgtC* when compared to the no treatment control (**Figure 1B**). When looking at c-di-GMP allosterically activated genes important for biofilm shown in our sequencing, bac7 treatment significantly decreased the expression of *bcsA* and *fimH* compared to the no treatment control (**Figure 1C**).

To understand the metabolic changes associated with bac7 treatment we quantified both ATP and c-di-GMP using the BacTiter-Glo microbial cell viability kit and cyclic di-GMP ELISA kit, respectively. Following 30-minutes treatment of hvKp NTUH-K2044, we found elevated ATP with bac7 treatment (**Figure 1D**). Control experiments using chloramphenicol (ribosomal biding control) and polymyxin B (lytic peptide control) treated samples demonstrated the expected increase and decreased in ATP, respectively (**Supplemental Figure 4A**). Conversely, we found significantly decreased c-di-GMP with bac7 treatment (**Figure 1E**). Control experiments revealed a similar decrease in c-di-GMP with chloramphenicol treatment, but not with polymyxin B (**Supplemental Figure 4B**). To assess if our transcriptional changes resulted in a phenotypical change, we quantified the abundance of FimA produced by hvKp NTUH-K2044 following treatment with increasing concentrations of bac7 compared to no treatment using FimA Immunoblot analysis (**Supplemental Figure 4C**). We found decreased FimA by immunoblot following treatment with 7.5 µmol L^−1^ bac7 (**Figure 1F**) and saw a significant reduction when quantified across several independent experiments (P=0.0067) (**Figure 1G**). Further, we performed transmission electron microscopy (TEM) and observed more bacterial cells expressing fimbriae in the control group compared to bacterial cells treated with 7.5 µmol L^−1^ bac7 (**Figure 1H; Supplemental Figure 4D**). Together our transcriptional analysis indicated bac7 causes both cytosolic and membrane stress, and we propose this dual stress causes an *mgtC*-mediated biofilm release by decreasing c-di-GMP and production of type I fimbriae.

### Bac7 transient membrane localization induces hvKp membrane depolarization but not leakage

Our RNA sequencing revealed bac7 increased the expression of genes encoding the membrane stress response protein RpoE and ribosomal subunits, together suggesting membrane and ribosomal stress are induced. Although bac7 has traditionally been described as a non-lytic peptide that kills via protein synthesis inhibition, Runti et al. described that this peptide has various modes of action depending on the bacterial genera ^24^. We hypothesize that although bac7 kills primarily via ribosomal inhibition, membrane stress is a contributing factor to the mechanism of killing and biofilm eradication in *K. pneumoniae*.

To determine if bac7 leaves the *K. pneumoniae* bacterial membrane intact similar *S. enterica* and *E. coli* bacterial membranes ^23^, we tested bac7 in a β-galactosidase leakage assay using o-nitrophenyl-β-d-galactopyranoside (ONPG) and measured the cytosolic pH using pHrodo™ dye to assess for leakage of hydrogen ions. We found no leakage (**Supplemental Figure 5A and 5B**) or changes in pH (**Supplemental Figure 5C**) with bac7 treatment confirming its non-lytic nature. We then assessed membrane depolarization with the hydrophobic cationic dye 3,3’-Dipropylthiadicarbocyanine Iodide (DiSC_3_) in a dose-response analysis over the course of 30-minutes, following a 30-minute pre-incubation with DiSC_3_ to allow for the dye to quench into the membrane. We found concentrations of 0.95, 1.9, and 3.8 µmol L^−1^ of bac7 depolarized the inner membrane of hvKp NTUH-K2044 within the first 10 minutes after the addition of the peptide (**Figure 2A**). This was then followed by a subsequent decrease in fluorescence intensity indicative of the re-quenching of the dye into the inner membrane. The fluorescence intensity of bac7 was not as high or as sustained as the positive control membrane lytic peptide cecropin A although well above the negative control non-lytic small molecule ertapenem (**Supplemental Figure 6**). We saw similar bac7 membrane depolarization with another hvKp strain KPPR1S (K2 capsule serotype) (**Figure 2B**). To take into account the role of capsular polysaccharides in the depolarization potential of bac7, we tested the capsular mutant KPPR1S *ΔwcaJ* ^41, 42^ that has an increased MIC (0.8 µmol L^−1^) compared to the parental isolate (0.24 µmol L^−1^) and found slightly more membrane depolarization at 1.9 and 3.8 µmol L^−1^. (**Figure 2C**). Similarly, when testing a colistin-resistant cKp strain MKP103, we found the membrane to have increased depolarization with bac7 treatment and minimal re-quenching (**Figure 2D**). Opposed to what we observed with hvKp NTUH-K2044, MKP103 showed slightly less depolarization with cecropin A lytic control than bac7 (**Supplemental Figure 6**). We next used high resolution single-cell fluorescence microscopy with FM-64 membrane dye and DAPI DNA stain to visualize the localization of a FITC N-terminal labeled bac7 peptide (FITC-bac7 MIC 1.9 µmol L^−1^) and found that with 15 minutes of incubation with 7.5 µmol L^−1^ of FITC-bac7, the peptide was enriched on the cellular envelope of hvKp NTUH-K2044 and KPPR1S, but not the KPPR1S *ΔwcaJ* capsular mutant (180 cl:3236 Figure 2C and **Supplemental Figure 7A**). Following 45 minutes incubation, we found, regardless of capsule abundance, the peptide had translocated into the cytosol. However, when testing bac7 at the MIC concentration (1.9 µmol L^−1^) we found at 15 minutes the peptide concentrated to the membrane of KPPR1S only (**Supplemental Figure 7B**). Transmission electron microscopy of hvKp NTUH-K2044 using a phosphotungstic negative stain to visualize the capsule and membrane following treatment with 7.5 µmol L^−1^ of bac7 revealed that although the membrane was largely intact, there was membrane blebbing and capsular changes (**Figure 2E**). Our results show that, although bac7 does not cause cytosolic leakage and pore formation like traditional host-defense peptides ^43^, its passage through the membrane to access the cytosol causes depolarization of the bacterial membrane leading to a membrane stress response and suggests a potential mechanism for the decrease in oxidative phosphorylation observed in our transcriptional analysis.

**Figure 2.**
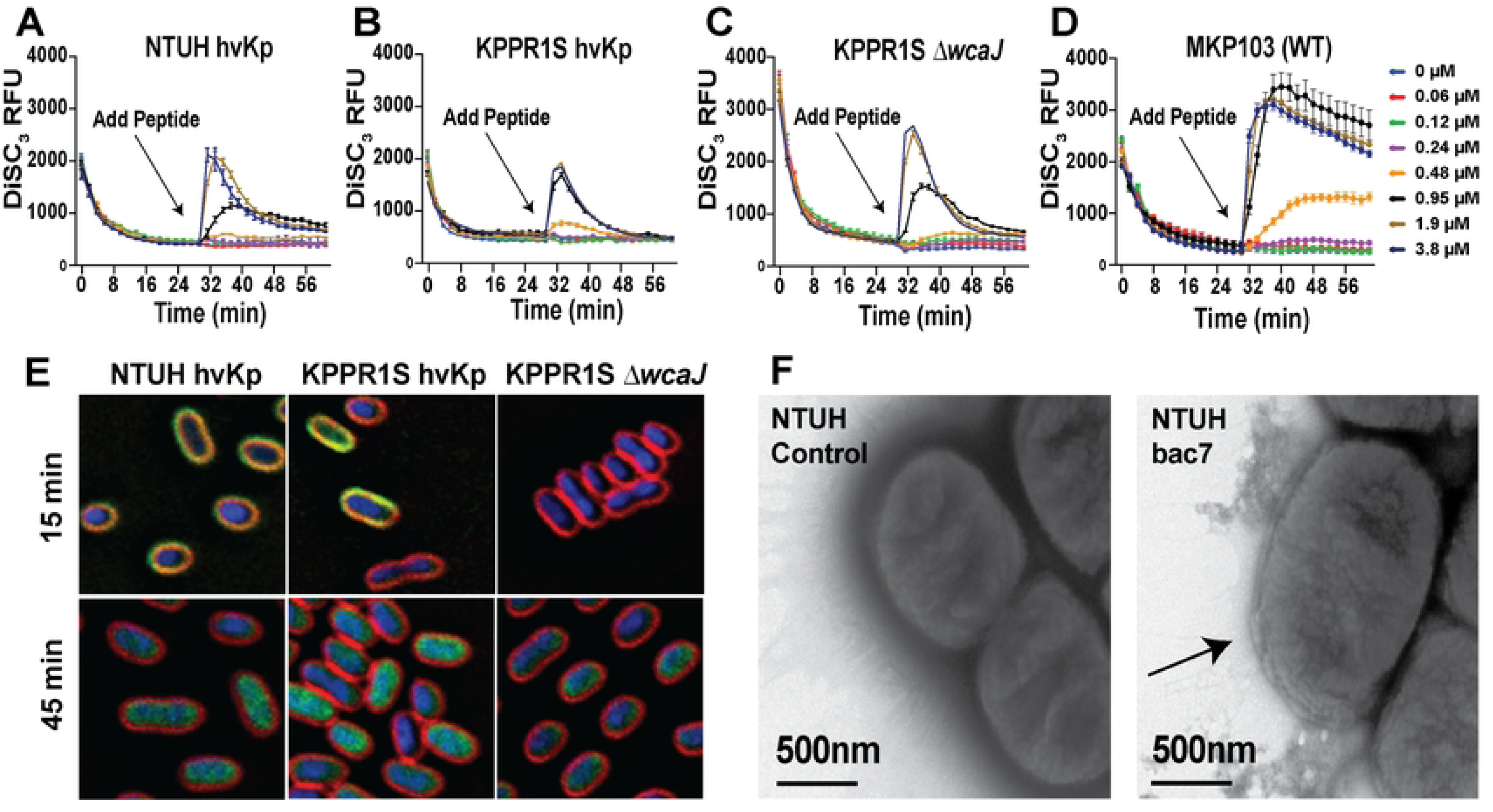
Bac7 transient membrane localization induces hvKp membrane depolarization but not leakage. Membrane depolarization of hvKp NTUH-K2044, hvKp KPPR1S, KPPR1S *ΔwcaJ,* and cKp MKP103 using a DiSC_3_ membrane potential dye (**Figure 2A-2D**). The dye was quenched into the membrane for 30-minutes before adding bac7 2-fold dilutions and the arrow indicates the peptide addition. Single cell fluorescent microscopy with FM 4-64 membrane dye (red), DAPI DNA stain (blue), after 15 minutes and 45 minutes incubation with 7.5 µmol L^−1^ FITC-bac7 (green) (**Figure 2E**). Transmission microscopy of hvKp NTUH-K2044 with a 0.1% phosphotungstic negative stain with or without 7.5 µmol L^−1^ bac7 (**Figure 2F**). Arrow indicates membrane blebbing locations with treatment.

### Bac7 displays differential membrane depolarization across *K. pneumoniae* clinical isolates

We hypothesize transient depolarization occurs as the peptide interacts with the membrane before entering into the cytosol as we observed in our fluorescence microscopy. Due to the variability between our hvKp and cKp lab strains (**Figure 2**), we wanted to expand our bac7 analysis beyond hvKp to broadly investigate membrane depolarization of clinical isolates from the *K. pneumoniae* MRSN diversity panel. For this, we performed DiSC_3_ membrane depolarization assays with bac7 as described with the lab strains.

We assessed the depolarization profiles of the MRSN isolates that displayed high mucoviscosity (N=17) and strong biofilm formation potential (N=15) in our previous work with these isolates ^27^. We found a diverse range of membrane depolarization of the MRSN isolates (**Supplemental Figures 8-13**). This diversity was not reserved to bac7 depolarization (**Supplemental Figures 8 and 11**) because we also found a diversity of membrane depolarization when we tested the lytic peptide cecropin A (**Supplemental Figures 9 and 12**), but as expected little depolarization with the beta lactam ertapenem (**Supplemental Figures 10 and 13**). To facilitate a broad comparison of the MRSN isolates, we graphed DiSC_3_ fluorescence increase with the lowest concentration that we see depolarization of the membrane (0.95 µmol L^−1^ bac7) at 10 minutes post-treatment, which is where we see peak fluorescence on the time course graphs, and at 30 minutes post-treatment to show the re-quenching of the dye (**Supplemental Figures 8-13**). Overall, we found that the mucoid isolates displayed increased membrane depolarization by bac7, as 12 isolates displayed increased depolarization compared to hvKp NTUH-K2044 (**Figure 3A; Supplemental Figure 14),** while only 8 strong biofilm formers displayed increased depolarization (**Figure 3B; Supplemental Figure 15**). Specifically, MRSN 365679 membrane exhibited more depolarization by bac7 than the cKp strain MKP103, while MRSN 1912 membrane was minimally depolarized. Interestingly, MRSN 21352 and 607210, are the two most mucoid isolates from our original study,^27^ and although this mucoidy is unlikely due to hyper-capsule as they lack the *rmpADC* operon that contributes to mucoidy ^7^, their membranes are very differently depolarized by bac7. To understand how the differential membrane depolarization of *K. pneumoniae* affects the killing kinetics of bac7 we performed a time-kill assessment at the respective 0, 0.5, 1, and 4X MICs (**Table 2**) of select MRSN isolates (MRSN 365679 and 1912) that displayed different levels of membrane depolarization. As expected from a protein synthesis inhibitor, we found bac7 to have bacteriostatic activity rather than bactericidal activity, although we did observe a shift to bactericidal activity after 24-hours with higher concentrations of bac7 towards MRSN 365679 (**Supplemental Figure 16**).

**Figure 3.**
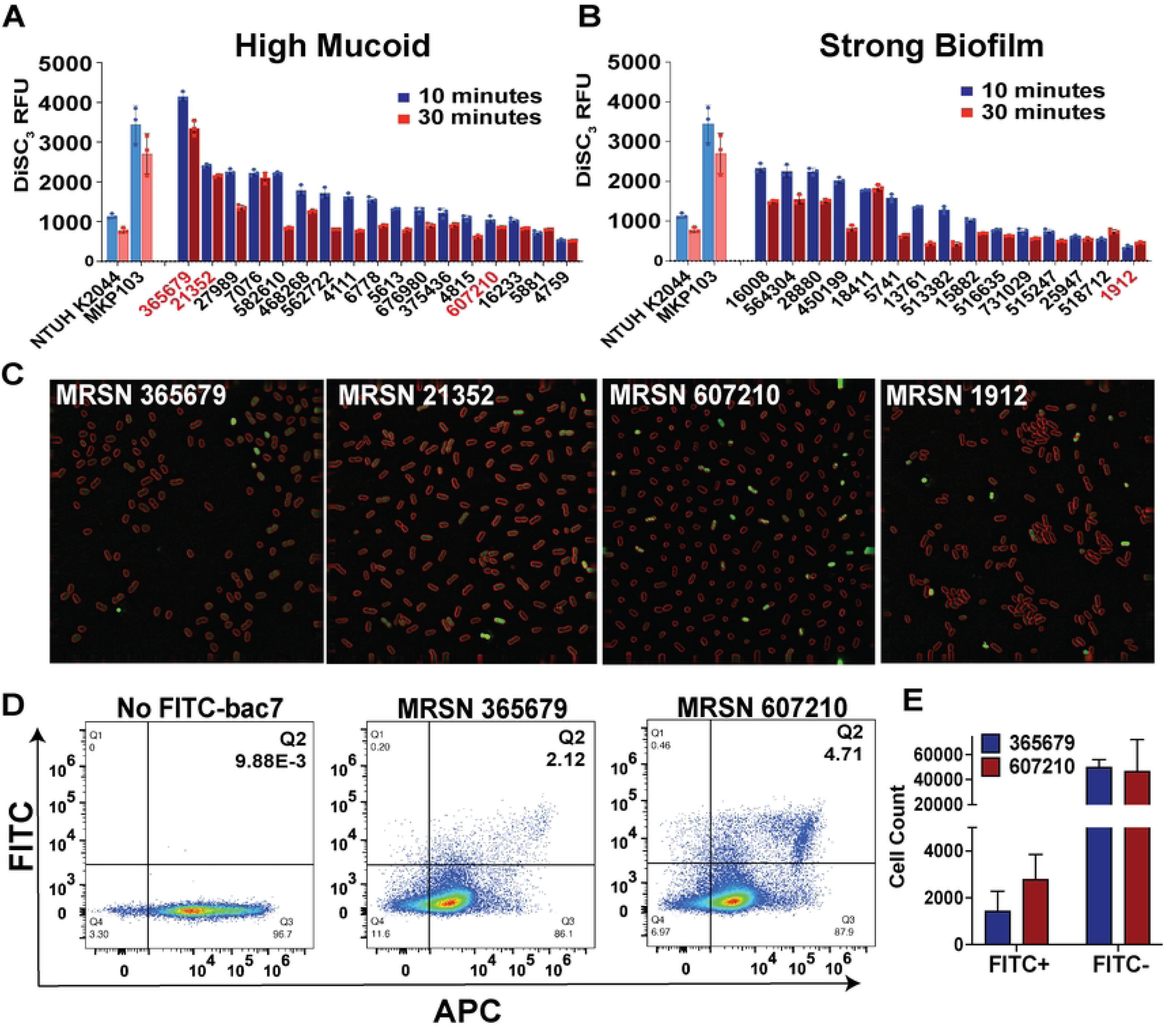
Bac7 displays differential membrane depolarization across *K. pneumoniae* clinical isolates. The figures show the membrane depolarization and FITC-bac7 uptake using the MRSN clinical diversity panel isolates. DiSC_3_ fluorescent readings after addition of 0.95 µmol L^−1^ bac7 for 10 minutes (peak fluorescence) and 30 minutes (quenching of fluorescence at endpoint) with the 15 most mucoid isolates (**Figure 3A**) and strongest biofilm forming isolates (**Figure 3B**) of the MRSN diversity panel. Control lab hvKp and cKp are shown for comparison in light blue (10 mins) and light red (30 mins). Isolates with red text in **Figures 3A** and **3B** were chosen for the single cell fluorescent microscopy. Single cell fluorescent microscopy with FM 4-64 membrane dye (red) after 15 minutes incubation with 1.9 µmol L^−1^ FITC-bac7 (green) (**Figure 3C**). Flow cytometry analysis with *Bac*Light™ red cell stain following 1.9 µmol L^−1^ FITC-bac7 treatment of MRSN 365679 and 607210 for 30 minutes using FITC and APC lasers for green and red fluorescence, respectively (**Figure 3D**). Gating was performed to generate 4 quadrants (Q1: FITC+/APC-; Q2: FITC+/APC+ Q3: FITC-/APC+; Q4: FITC-/APC-) with a high bar set to quantify super FITC fluorescent cells. Quantification of the cells in quadrant 2 (FITC+/APC+) and quadrant 3 (FITC-/APC+) with triplicate flow cytometry samples (**Figure 3E**). Single flow cytometry figures shown as representative of triplicate experiments in **Figure 3D**. Error for **Figures 3A**, **3B**, and **3E** are reported as ± SEM.

**Table 2.**
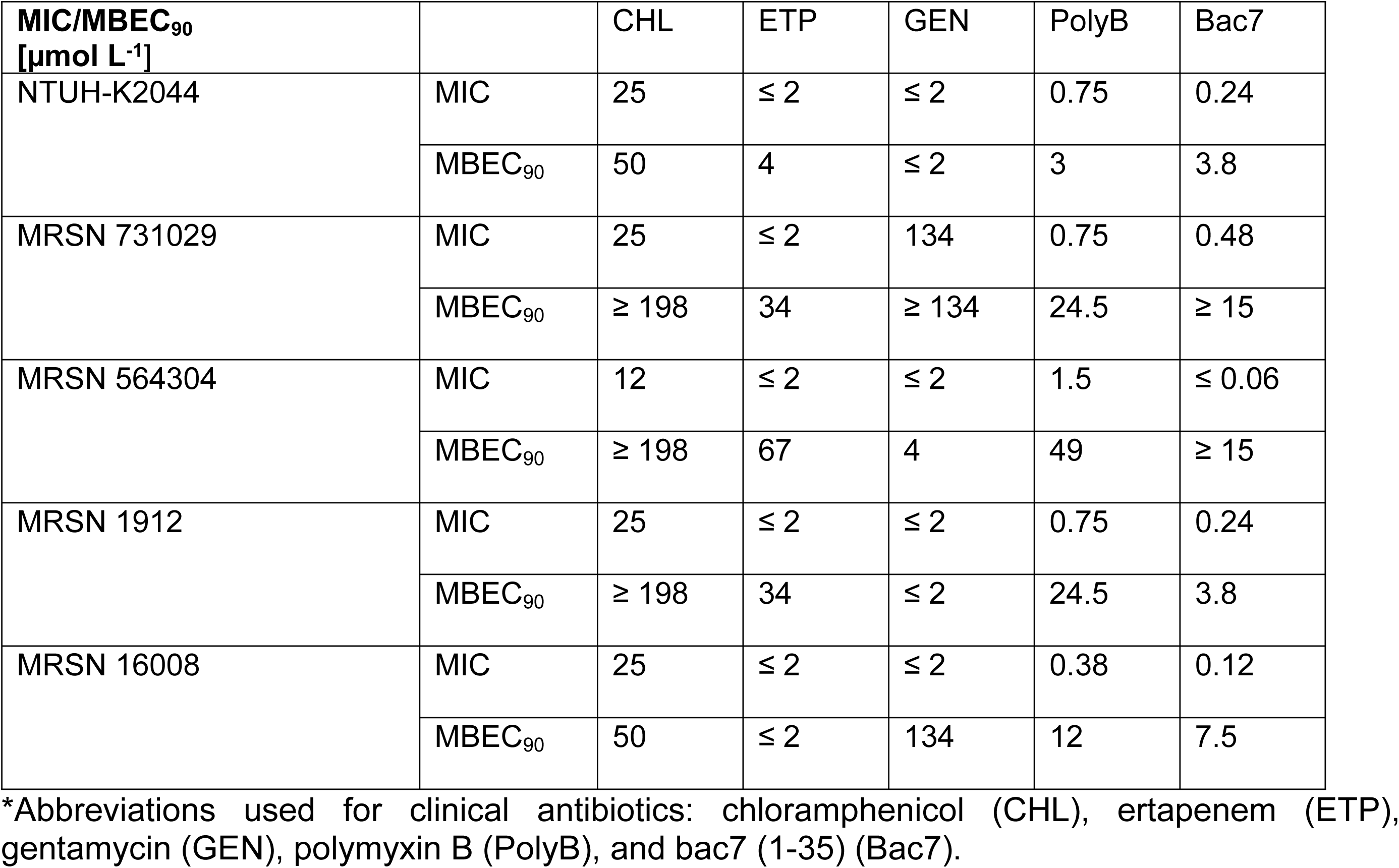
MIC and MBEC_90_ values of clinical antibiotics and bac7 towards *K. pneumoniae* isolates.

| MIC/MBEC<sub>90</sub><br>[μmol L<sup>-1</sup>] |  | CHL | ETP | GEN | PolyB | Bac7 |
| --- | --- | --- | --- | --- | --- | --- |
| NTUH-K2044 | MIC | 25 | ≤ 2 | ≤ 2 | 0.75 | 0.24 |
|  | MBEC<sub>90</sub> | 50 | 4 | ≤ 2 | 3 | 3.8 |
| MRSN 731029 | MIC | 25 | ≤ 2 | 134 | 0.75 | 0.48 |
|  | MBEC<sub>90</sub> | ≥ 198 | 34 | ≥ 134 | 24.5 | ≥ 15 |
| MRSN 564304 | MIC | 12 | ≤ 2 | ≤ 2 | 1.5 | ≤ 0.06 |
|  | MBEC<sub>90</sub> | ≥ 198 | 67 | 4 | 49 | ≥ 15 |
| MRSN 1912 | MIC | 25 | ≤ 2 | ≤ 2 | 0.75 | 0.24 |
|  | MBEC<sub>90</sub> | ≥ 198 | 34 | ≤ 2 | 24.5 | 3.8 |
| MRSN 16008 | MIC | 25 | ≤ 2 | ≤ 2 | 0.38 | 0.12 |
|  | MBEC<sub>90</sub> | 50 | ≤ 2 | 134 | 12 | 7.5 |

\*Abbreviations used for clinical antibiotics: chloramphenicol (CHL), ertapenem (ETP), gentamycin (GEN), polymyxin B (PolyB), and bac7 (1-35) (Bac7).

We then used isolates displaying either high (MRSN 365679 and 21352) or low (MRSN 607210 and 1912) membrane depolarization to test localization of a FITC-bac7 peptide after 15 minutes at 1.9 µmol L^−1^ using high resolution single-cell fluorescence microscopy as done with our hvKp isolates in **Figure 2E**. Although we did not see the same enrichment of FITC-bac7 to the cell envelope that we observed with our hvKp strains, we found a heterogenous localization pattern where a few cells displayed a high intracellular concentration of FITC-bac7 (**Figure 3C**). Interestingly, MRSN 365679 and 21352 displayed increased membrane depolarization but had fewer cells with concentrated peptide, compared to MRSN 607210 and 1912 that had less membrane depolarization by bac7 but more cells with concentrated peptide. Furthermore, cells with dense concentrations of peptide show cell rounding or have condensation of the cytosolic components. To understand these findings at a population level we performed flow cytometry analysis with MRSN 365679 (high depolarization/low uptake) and 607210 (low depolarization/high uptake) using *Bac*Light™ red cell stain following treatment with 1.9 µmol L^−1^ FITC-bac7 for 30 minutes. In line with our fluorescence microscopy imaging results, we found compared to MRSN 607210, MRSN 365679 had less cells with an extreme abundance of FITC-bac7 uptake (**Figure 3D** and **3E**). Overall, we found a strain-dependent membrane depolarization effect by bac7 revealing an inverse correlation between membrane penetration and depolarization of the bacterial membrane.

### Spatial distribution and cellular density generate biofilm mediated resistance in clinical isolate MRSN 564304

Our previous work revealed the potential of bac7 to collapse pre-formed biofilms of hvKp NTUH-K2044 ^26^. To expand on this to understand the broad effects of this peptide towards isolates with extremely robust biofilm formation capabilities, we assessed the anti-biofilm capabilities of bac7 towards four MRSN clinical isolates that have strong biofilm formation potential ^27^. We compared minimal inhibitory concentrations (MICs) to biofilm eradication of hvKp NTUH-K2044 and the MRSN isolates to measure the resistance attributed to biofilm formation. For this we used Calgary device plates for biofilm formation with crystal violet staining to define 90% minimal biofilm eradication concentration (MBEC_90_) of bac7 and clinical antibiotics. We found varying levels of biofilm mediated resistance (low MICs and high MBEC_90_s) (**Table 2**) with all clinical isolates displaying an in increased fold change in biofilm mediated resistance compared to NTUH-K2044 (**Supplemental Figure 17A**).

Our results show strong biofilm mediated resistance to bac7 treatment with MRSN 564304 (**Table 2**; **Supplemental Figure 17B**). To continue to investigate the resistance displayed by MRSN 564304 compared to the other isolates tested, we assessed the biofilm matrix spatial distribution of the four MRSN isolates using confocal z-stack imaging with SYTO 9 cellular stain (green) and calcofluor white polysaccharide stain (blue) to visualize the 3D spatial distribution of the cellular and polysaccharide populations, respectively ^27^. This technique is unique from crystal violet staining due to the ability to facilitate measuring of biofilm height and differentiation of the cells from matrix material. We found that 15 µmol L^−1^ of bac7 was able to collapse MRSN 1912 and 16008 (**Supplemental Figure 17C-17D**) more effectively than MRSN 731029 (**Supplemental Figure 17E**) and 564304 (**Figure 4A**). We then tested the potential for bac7 adjuvant therapy with the protein synthesis inhibitor chloramphenicol that binds to the 50S ribosomal subunit and is impacted by biofilm mediated resistance (**Supplemental Figure 17A**; **Table 2**) ^44^. We found that although bac7 or chloramphenicol (198 µmol L^−1^) alone were not effective against MRSN 564304 biofilm, when combined they effectively collapse this extremely robust biofilm (**Figure 4A**). We also saw combined collapse when testing this combination treatment against MRSN 731029 (**Supplemental Figure 17E**).

**Figure 4.**
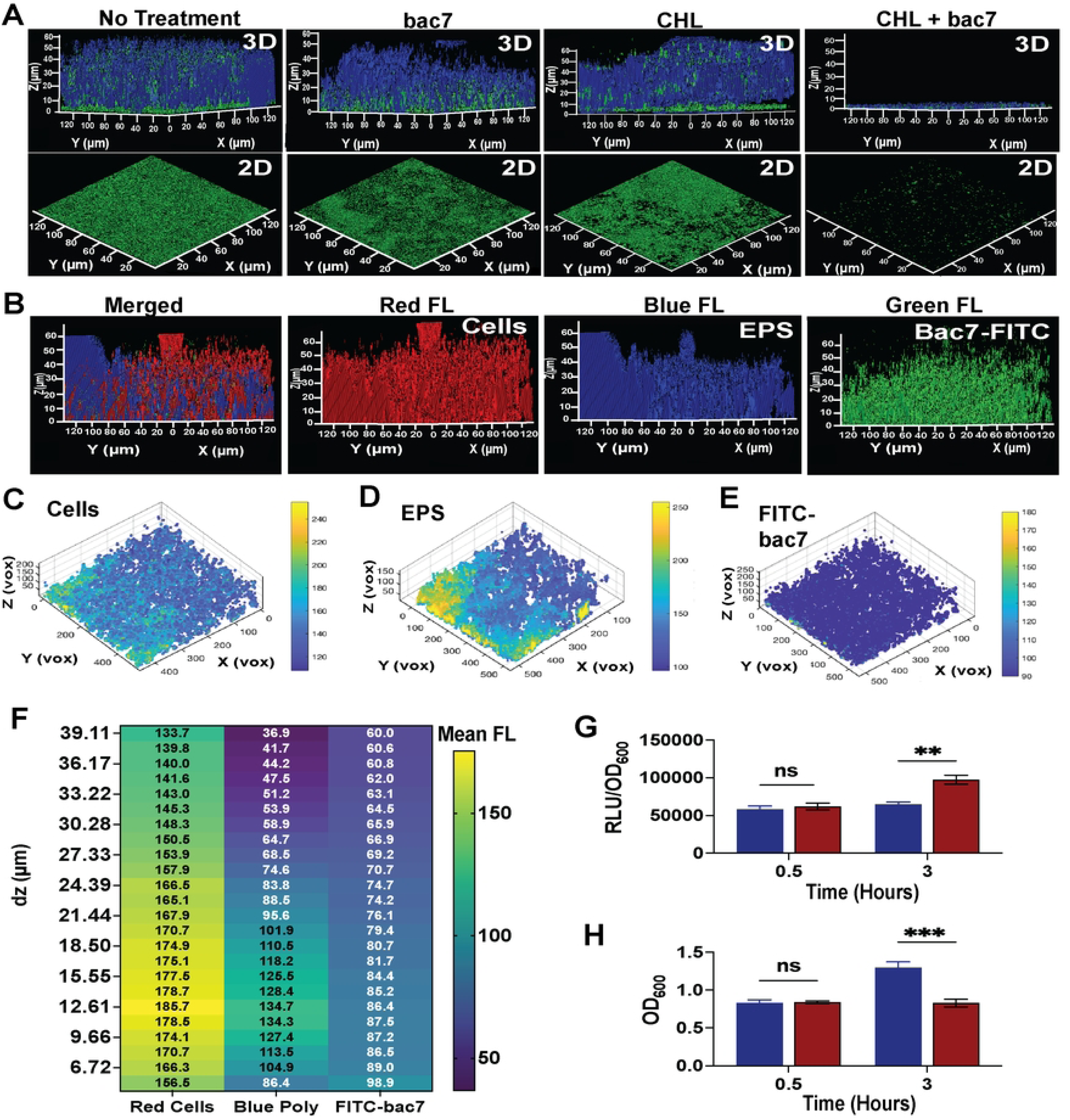
Spatial distribution and cellular density generate biofilm mediated resistance in clinical isolate MRSN 564304. The figures show confocal z-stack images of MRSN 564304 biofilm that is resistant to bac7 disruption. **Figure 4A** shows 3D and 2D images of confocal z-stacking imaging of MRSN 564304 biofilms using SYTO 9 dye cell stain (green) and calcofluor white polysaccharides (blue) following no treatment, bac7 (15 μmol L^−1^), chloramphenicol (CHL) (198 μmol L^−1^) or combined (bac7 +CHL). **Figure 3B** shows one-hour treatment of MRSN 564304 with 15 μmol L^−1^ FITC-bac7, stained with BactoView™ cell stain (red) and calcofluor white polysaccharide stain (blue) to visualize localization of the FITC-bac7 peptide (green). BiofilmQ software was used to process the confocal z-stack images and graph the data as 4D XYZC scatterplot to visualize the spatial quantification of the red cell stain (**Figure 4C**), blue polysaccharide stain (**Figure 4D**) and green FITC-bac7 peptide (**Figures 4E**). **Figure 4F** is a heat map of the mean relative fluorescence in each z-stack layer (dz (µm)) for all fluorophores. ATP quantification of MRSN 564304 biofilm embedded cells using relative luminescence units (RLU) normalized to the respective OD_600_ (RLU/OD_600_) using a CellTiter-Glo^®^ kit with no treatment and 15 µmol L^−1^ bac7 for 0.5 and 3 hours (**Figure 4G**). Biofilm OD_600_ MRSN 564304 for each timepoint used to normalize ATP RLU (**Figure 4H**). All imaging was performed in triplicate with one representative image shown. One-way ANOVA was used to determine significance compared to the no treatment groups for **Figures 4G** and **4H** with Dunnett’s correction for multiple comparisons and adjusted p-values shown (asterisks indicate p-values **** <0.0001, ** <0.01, and ns >0.1) and error shown reported as ± SEM.

We then treated biofilms of MRSN isolates with FITC-bac7 using confocal z-stack imaging with BactoView™ red nucleic acid stain (red bacterial cells) and calcofluor white dye (blue polysaccharide matrix), to visualize the localization of the FITC-bac7 peptide (green peptide). With one-hour treatment FITC-bac7 could not only penetrate the biofilm of a sensitive isolate MRSN 1912 (**Supplemental Figure 18A**), but also the resistant biofilm of MRSN 564304 (**Figure 4B**). When assessing 24-hour treatment of hvKp NTUH-K2044 and clinical isolates (MRSN 564304 and 1912), this validated that 15 µmol L^−1^ FITC-bac7 could penetrate all biofilms regardless of its anti-biofilm activity towards the isolate (**Supplemental Figures 19-21**). To understand the role of cell density of the resistant biofilms compared to the sensitive biofilms we quantified the cellular population and peptide density using biofilmQ image processing tool ^45^. This software tool is used to segment the confocal z-stack images to allow for spatial fluorescence quantification. We found that the mean intensity of fluorescence for the cells (**Figure 4C**) and matrix (**Figure 4D**) of the biofilm of MRSN 564304 were much greater than FITC-bac7 spatial fluorescence (**Figure 4E**). When comparing the mean fluorescence from the base to the top of the biofilms we observed FITC-bac7 fluorescence was higher throughout the layers of the matrix with MRSN 1912 (**Supplemental Figure 18B)** than with MRSN 564304 indicating the relative peptide concentration does not cover the cell density of this robust biofilm (**Figure 4F**). To understand if the cells within the biofilm of MRSN 564304 were being affected by bac7 we used the levels of ATP in the MRSN 564304 biofilm growing cells as a metric of peptide reaching its cytosolic target. Without treatment, MRSN 564304 displayed elevated intracellular ATP in biofilm embedded cells and increased biofilm density compared to MRSN 1912 (**Supplemental Figure 18C** and **18D**). We then treated pre-formed MRSN 564304 biofilms with 15 µmol L^−1^ bac7 and assessed ATP levels of the cells within the biofilm after 30 minutes and three hours. We saw a significant increase in ATP three hours post-treatment (**Figure 4G**) that aligned with a significant decrease in biofilm density compared to the no treatment group (**Figure 4H**). These results suggest that, although bac7 can penetrate the biofilm matrix of MRSN 564304 within one-hour, increased abundance of cells and elevated ATP within the biofilm-residing cells before treatment is likely the cause of the biofilm mediated resistance of this clinical isolate.

### Mutant analysis reveals both SbmA and MgtC are important for mitigating the biofilm disruption potential of bac7

We next aimed to understand more about bac7 membrane interactions and the consequences of triggering *mgtC* using bacterial mutants lacking the SbmA transporter and MgtC, respectively. We hypothesized that a mutant lacking SbmA would show increased depolarization as the peptide will be only accessing the cytosol by membrane penetration. Furthermore, we hypothesized from *S. enterica* literature ^38, 39^, that without MgtC there would be elevated ATP levels. We used transposon mutants of cKp MKP103 parental strain to test our hypotheses because of the high membrane depolarization by bac7 (**Figure 2D**), increased baseline expression of *phoP* and *mgtC* in normal growth conditions compared to hvKp NTUH-K2044 and KPPR1S (**Figure 5A**), and its extreme colistin resistance ^46^.

**Figure 5.**
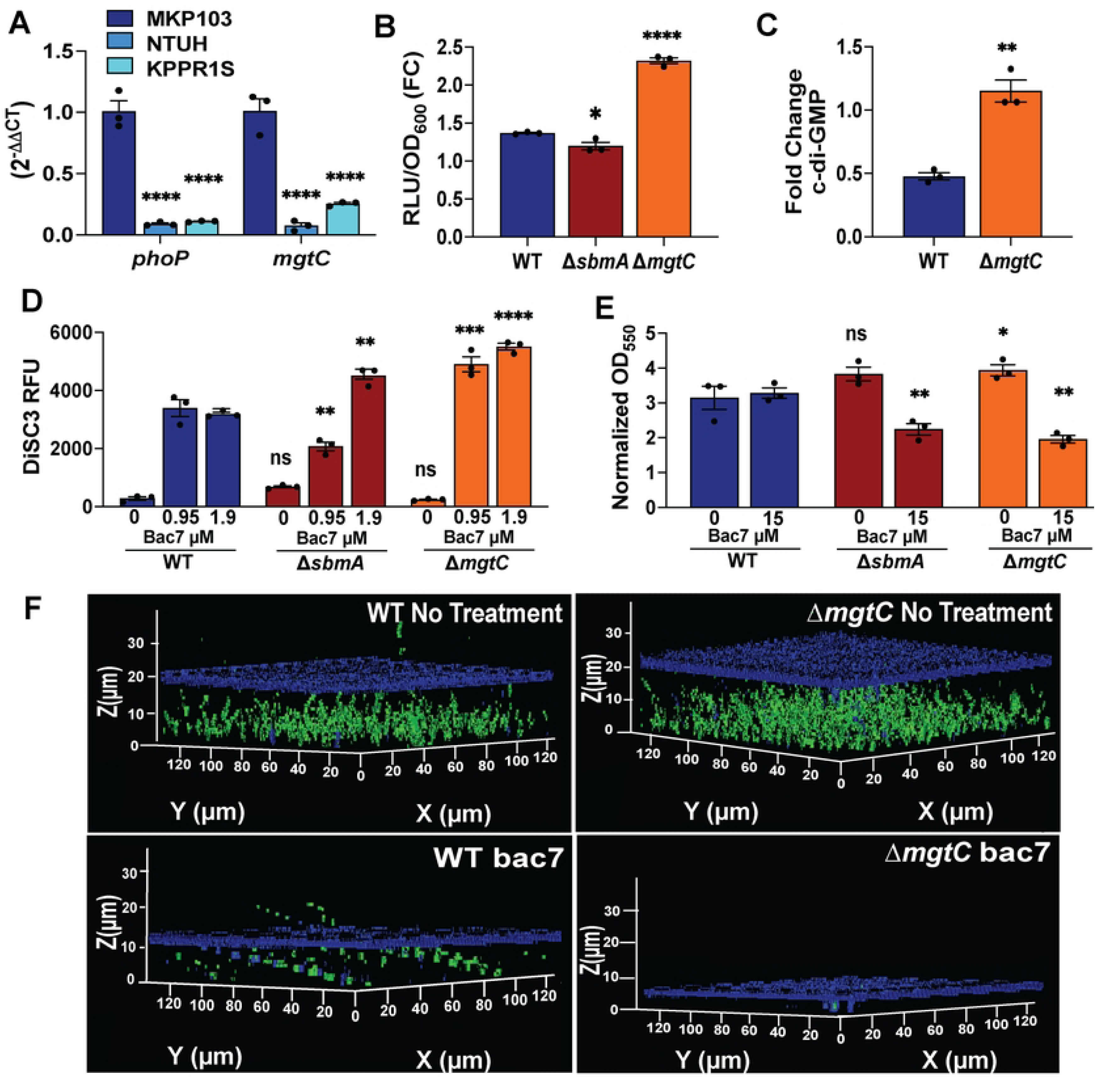
Mutant analysis reveals both SbmA and MgtC are important for mitigating the biofilm disruption potential of bac7. The figures show comparisons of the parental MKP103 (WT) and mutants lacking the SbmA polyproline transporter (*ΔsbmA*) and MgtC *(ΔmgtC).* Relative gene expression of *phoP* and *mgtC* grown in LB to log phase to analyze baseline expression of these genes in colistin-resistant MKP103 compared to hvKp NTUH-K2044 and hvKp KPPR1S using a 16S housekeeping gene for normalization (**Figure 5A**). Intracellular ATP levels shown as fold change in RLU/OD_600_ between no treatment and 1.9 µmol L^−1^ bac7 treatment for 30 minutes for parental MKP103 (WT), MKP103 *ΔsbmA*, and MKP103 *ΔmgtC* (**Figure 5B**). Fold change intracellular c-di-GMP levels between no treatment and 7.5 µmol L^−1^ bac7 for MKP103 parental (WT) and MKP103 *ΔmgtC* quantified using the cyclic-di-GMP ELISA kit (**Figure 5C**). DiSC_3_ membrane potential dye assessment of inner membrane depolarization with 0, 0.95, and 1.9 µmol L^−1^ bac7 showing the peak fluorescence observed following 10 minutes treatment for parental MKP103 (WT), MKP103 *ΔsbmA*, and MKP103 *ΔmgtC* (**Figure 5D**). Biofilm density measurement using crystal violet staining showing normalized (subtract background) biofilm density (OD_550_) after 24 hours with no treatment or 15 µmol L^−1^ bac7 (**Figure 5E**). 3D-rendering of confocal z-stack images of pre-formed biofilms of the MKP103 parental isolate (WT) and *ΔmgtC* mutant untreated and treated with 15 µmol L^−1^ bac7 (**Figure 5F**). Cells were stained with SYTO9 (green cells), and matrix polysaccharide were stained with calcofluor white (blue matrix). n=3 biofilms imaged for each condition with a representative image shown. Two-way ANOVA was used to determine significance for **Figures 5A** in comparison to MKP103 with Tukey’s multiple comparison correction, and one-way ANOVA was used to determine significance for **Figures 5C-5F** comparing the mutants to the WT with Tukey’s multiple comparison correction. Significance is shown with asterisks (adjusted p-values **** <0.0001, *** <0.001, ** <0.01, * <0.1, and ns >0.1) and error shown as ± SEM.

When assessing ATP levels in cKp MKP103, interestingly we do not observe the same increased intracellular ATP after 30 minutes with 1.9 µmol L^−1^ bac7 that we observed with hvKp NTUH-K2044, but found with this colistin-resistant strain that the RLU/OD_600_ closely matched the no treatment group (1.36-Fold Change) (**Figure 5B**). When comparing the mutant ATP levels to the parental strain (MIC 0.48 µmol L^−1^), we observed decreased intracellular ATP with *ΔsbmA* (KPNIH1_05310-803::T30) (MIC 3.8 µmol L^−1^) and increased intracellular ATP with *ΔmgtC* (KPNIH1_13815-409::T30) (MIC 0.48 µmol L^−1^). Intriguingly, we found even with no treatment we found increased intracellular ATP with *ΔmgtC* transposon mutant compared to the parental isolate (**Supplemental Figure 22A**). Similar to what we observed with the hvKp NTUH-K2044 treated samples, when assessing levels of c-di-GMP with the parental cKp MKP103 following treatment with bac7, there was a decrease in c-di-GMP compared to the no treatment, but increased c-di-GMP with the *ΔmgtC* transposon mutant (**Figure 5C**), which is similar to what we see with the ribosomal binding control tetracycline (**Supplemental Figures 22B** and **22C**). We then tested the membrane depolarization abilities of bac7 against the *ΔsbmA* and *ΔmgtC* transposon mutants (**Supplemental Figure 22D-22G**). We found a concentration dependent change in the peak fluorescence intensity (10 minute time point) with *ΔsbmA,* where with 0.95 µmol L^−1^ bac7 there was less depolarization of the membrane but more depolarization with 1.90 µmol L^−1^ bac7 compared to the parental strain (**Figure 5D**) that correlates with the loss of ATP observed with this mutant with bac7 treatment at this concentration. When comparing *ΔmgtC* transposon mutant to the parental strain there was an overall increase in membrane depolarization, where the lowest concentration to cause depolarization was 0.12 µmol L^−1^ (**Supplemental Figure 22D**) compared to 0.48 µmol L^−1^ with the parental strain (**Figure 2D**).

We then tested bac7 biofilm eradication at 15 µmol L^−1^ using crystal violet staining of MKP103 parental strain next to *ΔsbmA* and *ΔmgtC* transposon mutants and found both mutants were more sensitive to bac7 compared to the parental strain (**Figure 5E**). To further investigate the *ΔmgtC* transposon mutant biofilms, we used confocal z-stack imaging with SYTO 9 and calcofluor white dyes to visualize the biofilm changes in the mutant with and without peptide treatment. We found that without treatment *ΔmgtC* had increased biofilm cell density compared to the parental strain (**Figure 5F**). However, in line with the crystal violet staining, there is increased disruption of the *ΔmgtC* biofilm compared to the parental strain MKP103. Collectively, these results suggest bac7 is dependent on the SbmA transporter at low concentrations and that MgtC is functioning to relieve toxic ATP levels.

### *In vivo* treatment decreases bacterial burden and dissemination in a skin abscess biofilm infection model

In order to assess the physiologic effects of bac7 on a clinically-relevant *K. pneumoniae* biofilm, we leveraged a model of hvKp skin abscess and dissemination. To enable biofilm formation prior to treatment, we waited five hours after bacterial inoculation before treatment (peptide or the positive control polymyxin B) administration. Mice with a pre-formed linear skin abrasion were infected with 1×10^4^ colony forming units per milliliter (CFU mL^−1^) of hvKp NTUH-K2044. Five hours post-infection, a single dose of bac7 (20 µmol L^−1^; 10× MIC) or polymyxin B (10 µmol L^−1^; 10× MIC) was administered to the infection site. Mice were monitored for four days, with skin tissue and organs (liver, spleen, and kidneys) harvested on days two and four to quantify bacterial burden (**Figure 6A**). Mouse weight was recorded as a surrogate for disease severity (**Figure 6B**) ^47^. Notably, only bac7 treatment resulted in weight gain over the course of infection. The polymyxin B treated mice lost weight compared to the no treatment control group, indicating the mice could be experiencing adverse side effects that have been documented with this antibiotic ^48^.

**Figure 6.**
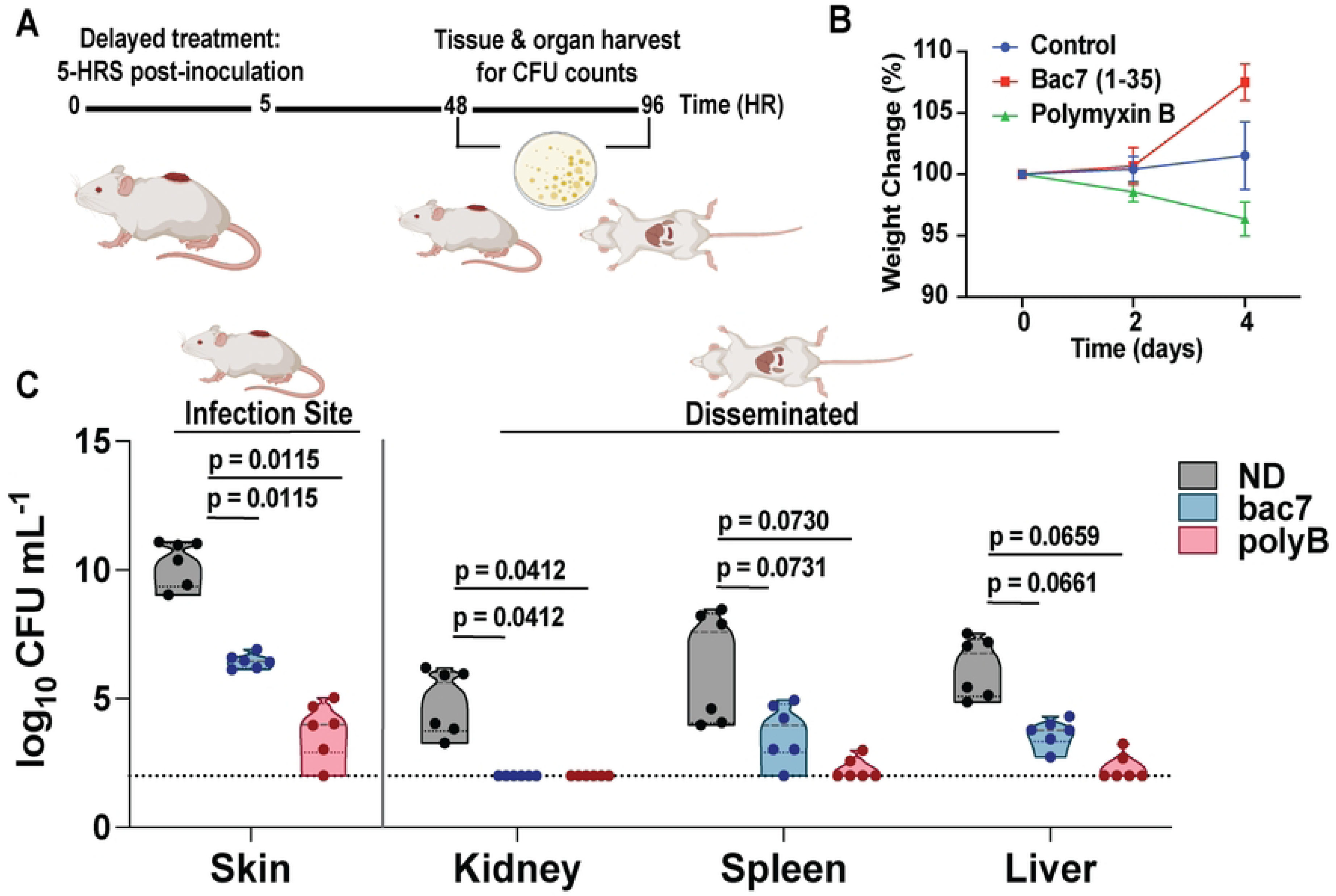
*In vivo* treatment decreases bacterial burden and dissemination in a skin abscess biofilm infection model. Skin abrasions on the ventral side of mice were formed and infected with 1×10^4^ CFU mL^−1^ hvKp NTUH-K2044. 5 hours post-infection, a single dose of bac7 and positive control polymyxin B was administered onto the infection site (**Figure 6A**). Four days post-treatment, regions of skin abrasions were excised, and organ systems were harvested and homogenized to quantify the bacteria. Mouse weight assessed at day two and day four post infection and shown as a percentage weight change compared to day 0 (**Figure 6B**). Log_10_ CFU mL^−1^ of the bacterial load in the skin, kidney, spleen, and liver (**Figure 6C**). Data was graphed as violin plots to show the mean values in addition to the upper and lower quartiles. Significance was determined using one-way ANOVA of n = 6 mice with p-values were obtained by comparing untreated groups to the treated groups. Biorender was used to generate the vector images with the figure designed in adobe illustrator.

At day two post-treatment, bacterial burden in skin tissues was reduced in both treatment groups compared to the untreated control (**Supplemental Figure 23**), although dissemination to distal organs was minimal at this early stage (10^2^-10^3^ CFU mL^−1^ compared to 10^6^ CFU mL^−1^ in skin tissue samples). In contrast, four days post-treatment, dissemination was more evident. Treatment with either bac7 or polymyxin B led to a significant reduction in CFU counts in the skin (p-value 0.0115) (**Figure 6C**). Bac7 also reduced bacterial load in the liver and kidneys. No bacteria were detected in the kidneys of either treated group at this time point. Spleen bacterial counts trended lower but did not reach statistical significance. Taken together, these results indicate that bac7 reduces local biofilm-associated bacterial burden, which limits dissemination of hvKp to distant organs.

## Discussion

Given the important contribution of biofilm formation to clinical infections and the extreme drug resistance observed in *K. pneumoniae*, targeting biofilms and understanding the underlying mechanisms of biofilm disruption are imperative steps towards better therapeutics. We previously described the biofilm disruption of hvKp NTUH-K2044 by bac7 and decreased mucoviscosity within the dispersed population ^49^. We found strong peptide aggregation due to a large portion of the N-termini bound to polysaccharides resulting in peptide-polysaccharide aggregation and biofilm collapse ^26^. Previous literature shows this peptide has different mechanisms of action between bacterial genera ^24^, but this had yet to be defined in *K. pneumoniae*. Here we characterize the mechanism of *K. pneumoniae* killing and biofilm disruption by bac7 and demonstrate a spectrum of biofilm disruption potential against a set of diverse clinical isolates. Our transcriptional data revealed oxidative phosphorylation was decreased in expression, ribosomal subunits are increased in expression, and changes in genes important for biofilm formation. Of note, we also found signs of membrane stress with this non-lytic peptide indicating it has a dual killing mechanisms, targeting both the membrane and ribosome. Intriguingly, we saw a gene encoding for an MgtC family protein was the second most increased in expression, and to date there are no studies into the function of this protein in *K. pneumoniae*.

The PhoPQ two component system facilitates a cascade of changes increasing outer membrane resistance to the last-resort antibiotic polymyxins ^50^. Within the PhoP regulon is *mgtC*, found to be important for controlling toxic levels of ATP experienced in the low pH and low magnesium intramacrophage environment ^38, 39^. However, ATP is a necessary component to produce c-di-GMP, an allosteric activator important for production of many important biofilm components and shown to be the molecular switch from sessile to motile phase in bacteria ^51^. Studies in *S. enterica* reveal MgtC not only decreases the production of cellulose but also decreases the bacterial membrane potential ^30,52^. MgtC has been shown to have a role in biofilm formation and exopolysaccharide production in *P. aeruginosa* ^53^. However, the function of this protein has yet to be determined in *K. pneumoniae*. We found that both *phoP* and *mgtC* had increased expression in our transcriptional data although there were extremely high levels of *mgtC* with peptide treatment. Within the *mgtC* upstream region hairpin autoregulation was shown to be induced by ATP ^40^. Therefore, we predict that once transcript of *mgtC* is produced through activation of the PhoP regulon, the elevated ATP levels caused by bac7 binding to the ribosomes is inducing the extreme increase in *mgtC* transcription. This resulted in decreased *bcs* and *fim* operon expression and decreased FimA production. We hypothesize that this is the cause of the rapid switch from biofilm formation before a significant change in viability.

Our previous work focused on the K1 capsule serotype hvKp NTUH-K2044, while here we expanded our analysis to test the potential biofilm disruption ability of bac7 toward a panel of clinical isolates. The MRSN diversity panel is a collection of clinical isolates recovered from the Walter Reed Army Institute of Research (WRAIR) from around the globe ^54^. When assessing the diversity of membrane depolarization caused by our bac7, we found mucoid isolates had overall higher depolarization. However, most of these isolates lack the *rmpADC* operon, indicating another factor is increasing mucoidy in these isolates ^27^. Furthermore, we saw a heterogeneity of peptide uptake, where some cells had an abundance of peptide while others had very minimal. This combined with the increase in expression of the genes encoding the VapBC toxin-antitoxin system suggest the presence of a subpopulation of persister cells resisting peptide uptake. Future work will be targeted at understanding the potential for resistance development towards polyproline peptides through persistence or sleeper cells. In addition, isolates with increased depolarization showed decreased peptide uptake when visualized via fluorescence microscopy and flow cytometry (**Figure 4**). We hypothesize that bac7 may be more slowly penetrating into the cells that display increased membrane depolarization. When testing biofilm resistance of the clinical isolates, we found MRSN 564304 had strong biofilm-mediated resistance to bac7. This isolate had elevated ATP and biofilm density compared to MRSN 1912. When looking at FITC-bac7 peptide penetration into biofilms, the confocal microscopy revealed penetration of the peptide into MRSN 564304 biofilm within one hour. However, using biofilmQ software we show the decreased peptide to cell ratio indicating insufficient cellular coverage.

Bac7 is in a class of proline rich peptides that enter the cell via the inner membrane protein transporter SbmA ^23^. However, some reports have shown membrane mechanism of action with this peptide in *P. aeruginosa* compared to *E. coli* ^28^. Our transcriptional analysis revealed there may be more than one mechanism in *K. pneumoniae*. Our single cell fluorescence imaging to visualize the entry of bac7 into the cytosol revealed transient membrane localization that we show leads to membrane depolarization and damage (**Figure 2**). Furthermore, we found diverse membrane depolarization abilities between different clinical isolates, with a potential correlation with decreased cell penetration and membrane depolarization (**Figure 4**). In the colistin-resistant cKp MKP103 this increased membrane depolarization correlated with less ATP increase following bac7 treatment, revealing increased depolarization aligns with decrease in peptide entry.

To provide a mechanistic insight into peptide penetration and role of MgtC in biofilm disruption we probed bacterial mutants lacking the SbmA transporter for peptide uptake and the MgtC like protein. We found bac7 does not induce the same levels of ATP in a colistin-resistant strain, but has elevated ATP in an *ΔmgtC* mutant, indicating MgtC is functioning as described in *S. enterica* ^38, 39^. When looking at the membrane depolarization potential we see more depolarization of the *ΔmgtC* mutant membrane at low concentrations but a concentration dependent increased depolarization with *ΔsbmA* where only at 1.90 µmol L^−1^ do we see increased depolarization compared to the parental strain. This shows bac7 is more dependent on the transporter at lower concentrations validating the increased MIC of this mutant (1.9 µmol L^−1^) while suggesting it is bypassing the membrane in a concentration dependent manner. Moreover, the increased depolarization of the *ΔmgtC* mutant shows increased membrane sensitivity in this mutant compared to the parental strain, suggesting that in *K. pneumoniae* MgtC may also function to control the membrane potential as has been shown in *S. enterica* ^52^. Future work will be focused on understanding the source and defining the consequences of the membrane depolarization induced by polyproline peptides. Finally, both mutants form a slightly stronger biofilm without peptide treatment compared to the parental strain but are more sensitive to bac7 treatment at 15 µmol L^−1^. This shows that peptide interactions with the membrane are important for biofilm disruption and suggesting that MgtC may actually play a role in protecting *K. pneumoniae* from bac7 biofilm disruption.

Finally, hvKp has emerged as a novel concern due to host immune evasion and dissemination of the bacterial infection to various organ systems through the bloodstream resulting in pyogenic liver abscess, endophthalmitis, renal abscess, and meningitis ^6^. With the changes in gene expression, we modeled a *K. pneumoniae* biofilm abscess infection. With the loss of matrix material and decreased mucoidy of the dispersed population observed in our previous work, we hypothesized that this would decrease dissemination of hvKp. We demonstrate the ability of bac7 to decrease the bacterial burden in the wound and to prevent dissemination from the site of infection to the kidneys, liver, and spleen (**Figure 6**). Intriguingly, the bac7 treated mice, as opposed to the polymyxin B mice, gained weight throughout the four-day experiment indicating the increased health with this treatment comparatively. This is an important factor considering the clinical toxicity shown by polymyxins ^48^. Although preliminary, these findings suggest the potential of bac7 to prevent dissemination from a wound. Future studies will focus on other routes of infection to determine the range of efficacy in a variety of environmental niches that influence the *K. pneumoniae* phenotype.

In conclusion, we have shown the dual action of bac7 can disrupt biofilms formed by hypervirulent and drug-resistant *K. pneumoniae* pathotypes, including clinical isolates. We reveal bac7 treatment potentially switches *K. pneumoniae* from biofilm state through regulation of cellulose and fimbriae. While this would be less advantageous for a non-antimicrobial, bac7 also imparts strong antibacterial activity towards the cells released from the biofilm for a unique combination of anti-biofilm and anti-bacterial mechanisms. Using a delayed treatment *in vivo* skin abscess model that emulates a biofilm mediated infection, we demonstrate that bac7 is a promising therapeutic for decreasing dissemination of biofilm-associated hvKp infections.

## Methods

### Bacterial strains

Bacterial lab strains used in this study are listed in **Supplemental Table 2.** The Walter Reed Army Institute of Research Multidrug-Resistant Organism Repository and Surveillance Network (MRSN) diversity panel of *K. pneumoniae* isolates were used for the clinical isolates in this study are listed in (**Supplemental Table 3**). All overnight cultures were grown in Luria-Bertani (LB) broth at 37°C with 220 rpm shaking. Overnight cultures were synchronized to log phase using 1:10 subculturing into LB broth and grown for three hours 37°C with 220 rpm shaking. Chloramphenicol 100 µg mL^−1^ was supplemented to LB when growing cKp MKP103 transposon mutants.

### Peptides

Bac7 (1-35) peptide was ordered from Novopro (novoprolabs.com/p/bac7) and polymyxin B sulfate from TCI chemicals (https://www.tcichemicals.com). Novopro synthesized the N-terminal FITC tagged bac7 peptide used for the microscopy and flow cytometry analyses. All peptides were resuspended in ultra-purified water at 10 mg mL^−1^ and stored at −20°C in 50 µL aliquots.

### Transcriptional analyses

#### Bac7 treatment and RNA extraction

Overnight cultures of hvKp NTUH-K2044 were synchronized for 3 hours diluted to an optical density 600 nm (OD_600_) 0.5 in Mueller Hinton Broth 1 (MHB1), treated with 7.5 µmol L^−1^ bac7 for 30 minutes in a water bath at 37 °C, in parallel with no treatment controls. After incubation, the samples where centrifuged at 15,000 × g and processed for total RNA extraction using the hot phenol procedure as previously described with modifications ^12^. Briefly, RNA was resuspended in 10 µl nuclease free water and quantified by Nanodrop. The RNA was then DNase treated using TURBO DNA-*free*™ Kit (AM1907, Invitrogen). RNA samples were aliquoted and stored at −80 until downstream analysis by RNA sequencing and RT qPCR.

#### RNA-sequencing

The RNA samples were submitted to SeqCenter (https://www.seqcenter.com/) and sequenced using a NovaSeq X Plus System. Read quantification was performed using Subread’s feature Counts functionality ^55^. After normalization, read counts were converted to counts per million (CPM). Differential expression analysis was performed using edgeR’s glmQLFTest. Differentially expressed genes (DEGs) were identified as the genes with logFC > 1 and FDR < 0.01 between the bac7 treated samples and no treatment controls. Pathway analysis was performed using limma’s “kegga” functionality ^56^. The genes that were considered Up/Down in this analysis were at FDR < 0.05.

#### Protein–protein interaction (PPI) network construction

A subgroup of annotated DEGs after a stringing filtering by LogFC and FDR (Log FC >1.5, FDR < 10e-10) for a total of 216 genes were uploaded to the Search Tools for the Retrieval of Interacting Genes (STRING, http://www.string-db.org/) ^57^. In the PPI network analysis, a confidence > 0.9 was defined as the cut-of criterion.

#### Reverse transcriptase quantitative PCR (RT-qPCR)

RNA samples were reverse transcribed using SuperScript™ IV VILO™ Master Mix (11756050, Invitrogen) following the manufacturer’s instructions. RT-qPCR was performed using PowerUp™ SYBR™ Green Master Mix (A25742, Applied Biosystems) following the manufacturer’s instructions, and the QuantStudio 3 Real-Time PCR System (A28567, Applied Biosystems). There were three technical replicates against three biological replicates for qPCR analysis. The primers used for RT-qPCR were designed using the Integrated DNA Technologies (IDT) PrimerQuest tool. Primer sets used are listed in **Supplemental Table 3**. Relative expression levels of the target transcripts were calculated using the 2^−ΔΔCt^ method ^58^. *ftsZ* and 16S rRNA were used as endogenous housekeeping genes. Two-way ANOVA was used to determine significance of the gene expression changes.

### Intracellular ATP quantification

#### Planktonic

Overnight cultures were grown in lysogeny broth (LB) media at 37°C in shaking incubator (220 rpm) for 24 hrs. Overnight cultures were synchronized 1:100 in LB media for 3 hours in shaking incubator at 37°C. Synchronized cultures were standardized to OD 0.5 in MHB1 media and 2 mL culture was pipetted into culture tubes with 30× MIC of the controls polymyxin B and chloramphenicol or 15 µmol L^−1^ bac7. Cultures were incubated in shaking incubator (220 RPM) at 37°C for 30 minutes. Cultures were centrifuged at 21,130 × g for 10 minutes and pellets were resuspended in 1 mL LB media. Samples were heat-killed in the 70°C water bath for 1 hr. 100 µL of samples were pipetted into a 96-well black-walled clear bottom plate (Corning^®^) with 100 µL CellTiter-Glo^®^ 2.0 Cell Assay reagent and placed into a BioTek SYNERGY H1 plate reader for 2 minutes of orbital shaking followed by 10 minute incubation. Luminescence was read after 10 minutes incubation. Results were measured in relative luminescence units (RLU) and normalized to the respective OD_600_ values. Fold change values were achieved by dividing RLU/OD_600_ of the treated samples by the RLU/OD_600_ of the no-treatment control samples and graphed with error reported as ± SEM.

#### Biofilm

Biofilms were grown and treated as described for the confocal imaging above and incubated in 37°C static incubator for 30 minutes or three hours. Dispersed cells and biofilms were collected into 1.5 mL microcentrifuge tubes and heat-killed at 70°C for 1 hour in water bath. 100 µL heat killed biofilm and dispersed cultures were analyzed as described above for the planktonic treatment cultures.

### c-di-GMP quantification

Quantification of c-di-GMP was performed using a Cyclic di-GMP ELISA (Cayman Chemical). Overnight cultures of hvKp NTUH-K2044, MKP103 WT, and MKP103 Δ*mgtC* were synchronized for three hours and treated with 7.5 µmol L^−1^ bac7 for 30 minutes at 37°C with 220 rpm shaking. The cultures were centrifuged at 15,000 × g and resuspended in Wash Buffer (#400062) with 0.5 µg mL^−1^ polysorbate. Samples were then sonicated on ice for 20 seconds with 59 seconds delay for a total of 15 minutes and centrifuged at 15,000 × g for 5 minutes at 4°C. The assay was then performed according to kit instructions (Caymanchem.com/product/501780/cyclic-di-gmp-elisa-kit). Briefly, Cyclic di-GMP ELISA Standards (#401784) were made by diluting the bulk standard 1:3 into Immunoassay Buffer C (1×) (#401703). Tracer dye (#401780) was made by adding 60 µL of dye into 6 mL tracer for a 1:100 final dilution. Antiserum Dye (#401782) was prepared by adding 60 µL of dye into 6 mL antiserum for a final dilution of 1:100. The wells of the provided 96-well plate were washed five times with 300 µL of Wash Buffer (1×). Plate layout was followed as suggested by ELISA kit. 100 µL Immunoassay Buffer C (1×) was added to Non-Specific Binding (NSB) wells and 50 µL was added to Maximum Binding (B_o_) wells. 50 µL of ELISA Standards were added to the respective wells. 50 µL samples were added to appropriate wells and 50 µL tracer was added to all wells except Total Activity (TA) and Blank (Blk) wells. 50 µL Antiserum was added to all wells except TA, NSB, and Blk wells. 96-well plate was covered and placed in BioTek SYNERGY H1 plate reader for 2 hours at room temperature with continuous orbital shaking. Wells were emptied and washed with 300 µL Wash Buffer (1×). 175 µL TMB Substrate Solution (#400074) was added to all wells and 5 µL tracer was added to TA wells. Plate was covered and placed in BioTek SYNERGY H1 plate reader for 30 minutes at room temperature with continuous orbital shaking. 75 µL of HRP Stop Solution (#10011355) was added to all wells. Plate was read in BioTek SYNERGY H1 plate reader at wavelength of 450 nm.

### Western blotting for type I fimbriae

Overnight cultures of hvKp NTUH-K2044 were synchronized for two hours to an OD_600_ between 0.4 and 0.6. Synchronized cultures were treated with 7.5 µmol L^−1^ bac7 for 30 minutes in a water bath at 37°C. Treatment with sterile 1× PBS was used as a negative control (e.g., 0 µmol L^−1^ bac7). Cultures were centrifuged, washed 1x with sterile 1× PBS, and resuspended in 50µl of PBS. 6× SDS and 1M HCL were added to samples that were then boiled at 95°C for 5 minutes. 1M NaOH was added to samples after cooling. After treatment, samples were run on SDS-PAGE gels. Gels were transferred to a 0.2 µm PVDF membrane using semi-dry transfer system. Transferred membranes were blocked overnight in 2%BSA and 3% milk in PBS. Membranes were washed (0.1% PBST) then stained with either 1:2,000 1°Rabbit anti-whole type I pili and 1:2,000 Rpo alpha subunit isolated from *E. coli* (loading control) for 2 hours rocking at room temperature. After 2hrs, membranes were washed and treated with 1:10,000 Goat anti-rabbit IgG HRP or 1:5,000 Sheep anti-mouse for 2 hours rocking at room temperature. Following treatment with 2° antibodies, membranes were washed and exposed with ECL (Bio-Rad) before imaging. Blots were analyzed using ImageJ software and GraphPad Prism v10. Mouse anti-RNA Polymerase (Biolegend cat. 663104) was used for RNA-polymerase control staining. Rabbit anti-Type 1 pilus antibody was produced by immunizing soluble pilus protein in rabbits by New England Peptide (now Biosynth, Louisville, KY). Whole pili were purified using the *E. coli* strain AAEC185 with the plasmid pBAD18-Cm-*fimA-H*, containing the pilus operon from *K. pneumoniae* TOP52. Cultures were grown overnight in LB at 37°C with 0.2% arabinose. Pili were purified by heating to 65°C, shearing via vortex, and salt precipitation with MgCl_2_ as previously described ^59^. The soluble pilus fraction was used for vaccination of rabbits and antibody preparation.

### Membrane depolarization assay

Bacterial cultures were grown overnight and synchronized as described in the bacterial killing assays. Following 3-hour synchronization, samples were washed three times with 15 mL Buffer A (5 mM HEPES, 5.0% glucose) and resuspended in 10 mL Buffer A. The resuspended cells were standardized to OD_600_ 0.1 in 25 mL Buffer A with 100 mM KCl and incubated in shaking incubator (150 rpm) at 30°C for 15 minutes. Following the brief incubation, 2 µmol L^−1^ DiSC_3_(5) (3,3’-Dipropylthiadicarbocyanine Iodide) (ThermoFisher) was added to the cell suspension and was pipetted into a 96-well black-walled clear bottom plate (Corning®). Fluorescence measurements (excitation 622 nm, emission 670 nm) were recorded every 2 minutes at 37°C using BioTek SYNERGY H1 plate reader to detect fluorophore quenching. Following the initial 30-minute quenching, 2-fold bac7 dilutions using buffer A were prepared alongside positive control peptide cecropin A (anaspec) and negative control ertapenem (Fisher Scientific) in triplicate as described in the minimal inhibitory concentration assay and 50 µL of the peptide dilutions were transferred to the black-walled clear bottom 96-well plate containing the cell suspension with quenched DiSC_3_ dye. Fluorescence measurements (excitation 622 nm, emission 670 nm) were recorded every 2 minutes at 37°C for an additional 30 minutes. The fluorescence readouts were graphed with error reported as ± SEM.

### Single cell fluorescence microscopy imaging

Cells were prepared and imaged as previously described ^60, 61^. Briefly, cultures were streaked out and grown overnight at 37°C. Cultures were then standardized to an OD_600_ of 0.1 in 2 ml in LB and FITC-bac7 was added. Some samples were incubated at 37°C shaking at 230 RPM for 30 minutes. 1 ml of culture was spun down and resuspended in 100 µl of supernatant. 5 µL of culture was spotted onto a glass bottom dish (Mattek, P35G-1.5-14-C) and cells were then stained with 1 μg mL^−1^ FM-464 fluorescent dye (MilliporeSigma, 574799-5MG) to visualize the membrane and covered with an agarose pad (a process that takes 15 minutes which is included in the incubation times). Images were captured on a DeltaVision Core microscope system (Leica Microsystems) equipped with a Photometrics CoolSnap HQ2 camera and an environmental chamber. Seventeen planes were acquired every 200 nm and the data was deconvolved using SoftWorx software. Images were created using Fiji ^62^.

### Transmission electron microscopy

#### Negative stain for membrane damage assessment

*K. pneumoniae* NTUH-K2044 was grown overnight and diluted to an OD_600_ 0.1 in Mueller Hinton Broth with 7.5 µmol L^−1^ bac7 and incubated for 15 minutes at 37°C. After peptide treatment for 15 minutes the media was removed, and the cells were resuspended in an equal volume of 0.2× PBS so that the salts did not interfere with imaging. Formvar-carbon coated grids (EMS) were glow discharged for one minute before adding 5 µL of sample and the sample was allowed to sit for 2 minutes. Grids were then blotted, washed with a drop of water, blotted again, and then stained with 5 µL of 1% phosphotungstic acid (PTA) (pH 7) for one minute. The stained samples were blotted and allowed to air dry before imaging using a Technai Spirit TEM at 80kV. Images used for the figures were taken at 26.5kx and 43.5kx magnification. The microscopy imaging was performed at the University of Texas Center for Biomedical Research Support (CBRS) core.

#### Fimbriae analysis

Strains were cultured as described above for Western blotting. NTUH-K2044 was grown overnight and synchronized to an OD_600_ between 0.4-0.6. Cultures were washed with 1× PBS and resuspended in 1× PBS. Samples were then prepped for negative-stain EM by fixing in 1% glutaraldehyde for 10 minutes. The bacterial suspension was allowed to absorb onto freshly glow discharged formvar/carbon-coated copper grids (200 mesh, Ted Pella Inc., Redding, CA) for 10 min. Grids were then washed two times in dH2O and stained with 1% aqueous uranyl acetate (Ted Pella Inc.) for 1 min. Excess liquid was gently wicked off and grids were allowed to air dry. Samples were viewed on a JEOL 1200EX transmission electron microscope (JEOL USA, Peabody, MA) equipped with an AMT 8-megapixel digital camera (Advanced Microscopy Techniques, Woburn, MA). Percent piliation was determined by manually counting at least 50 cells/sample and dividing the number of cells with extruding pili by the total number of cells counted.

### Inner membrane permeability assay

Bacterial cultures were grown overnight and synchronized as described in the bacterial killing assays. Following synchronization, the bacteria were centrifuged and washed with 25 mL of 100 mM sodium phosphate buffer. Centrifugation was repeated and bacteria was resuspended in 5 mL 100 mM sodium phosphate buffer and OD_600_ reading was recorded. Bac7 was diluted alongside positive control peptide cecropin A and negative control ertapenem using 2-fold dilutions in 100 mM sodium phosphate buffer in a 96-well black-walled clear bottom plate (Corning®). Culture was standardized to OD_600_ 0.1 in 100 mM sodium phosphate buffer with 1.5 mM ONPG (o-nitrophenyl-β-D-galactopyranoside) was added to all wells. Fluorescence measurements (excitation 622 nm, emission 670 nm) were recorded every 2 minutes for 45 minutes using a BioTek SYNERGY H1 plate reader. The fluorescence readouts were graphed with error reported as ± SEM.

### Intracellular pH assessment

Bacterial cultures were grown overnight and synchronized as described in the bacterial killing assays. Following synchronization, 50 mL culture was centrifuged at 15,000 × g for 10 minutes at room temperature. Supernatant was removed and bacterial pellet was resuspended in 10 mL 5 mM HEPES buffer with 5 mM glucose. Centrifugation was repeated with the same parameters above. Supernatant was removed and pellet was resuspended in 10 mL 5 mM HEPES buffer with 5 mM glucose. OD_600_ reading was recorded, and culture was standardized to OD_600_ 0.1 in 5 mM HEPES buffer with 5 mM glucose. 2-fold serial dilutions of drug were performed in 96-well black-walled clear bottom plate (Corning®). 50 µL of standardized culture was added to wells of 96-well black-walled clear bottom plate (Corning®) containing drugs and was incubated in 37°C static incubator for 30 minutes. 12 µmol L^−1^ pHrodo™ red fluorescent dye (Invitrogen) was prepared in 5 mM HEPES buffer with 5 mM glucose. 100 µL of 12 µmol L^−1^ pHrodo™ red fluorescent dye (Invitrogen) was added to wells of 96-well black-walled clear bottom plate (Corning®) containing bac7, chloramphenicol, and polymyxin B and was incubated for 30 minutes at room temperature to allow for uptake of dye. Absorbance was read at 533 nm fluorescence reading using a BioTek SYNERGY H1 plate reader. The fluorescence readouts for n=3 replicates were graphed with error reported as ± SEM.

### Flow cytometry with FITC-bac7

To understand the heterogenicity of peptide uptake with a large population of cells we used flow cytometry following treatment with 1.9 µmol L^−1^ FITC-bac7 for 30 minutes. Following treatment, the cells were centrifuged to remove remaining peptide and resuspended in phosphate buffered saline with 200 µmol L^−1^ *Bac*Light™ Red Bacterial Stain (Invitrogen) and incubated in the dark at room temperature for 15 minutes. A CytoFlex flow cytometer was used to analyze the samples using FITC laser and APC red fluorescent laser. The no peptide control was used to gate for no green fluorescence before analyzing the samples with FITC-bac7 treatment. FlowJo software was used to generate graphs used for the figures shown. The samples were run in triplicate and the total count of FITC+/Red+ and FITC-/Red+ for all three samples were graphed with error shown as ± SEM.

### Bacterial killing assays

#### Minimal inhibitory concentration assays

Overnight cultures were grown in lysogeny broth (LB) media at 37°C in shaking incubator (220 rpm) for 24 hours. Overnight cultures were synchronized 1:100 in LB media at 37°C in shaking incubator (220 rpm) for 3 hours and standardized to an OD_600_ 0.002 in Mueller Hinton Broth 1 (MHB1) media. Bac7 stock concentration (10 mg mL^−1^) was diluted in bovine serum albumin (BSA) buffer (BSA and 0.1% acetic acid) to 32 µg mL^−1^ in triplicate and diluted 2-fold in 96-well plate using BSA. Following drug dilution, 50 µL standardized culture was added to all wells containing 50 µL of the drug dilution. The 96-well plate was sealed using Parafilm and placed in 37°C static incubator for 24 hours. 96-well plate was read at OD_600_ after 24 hours. For minimum inhibitory concentration assays testing clinical antibiotics, the methods detailed above were utilized with minor modifications. Overnight cultures were synchronized 1:100 in LB media at 37°C in shaking incubator (220 rpm) for 3 hours and standardized to an OD_600_ 0.002 in Mueller Hinton Broth 2 (MHB2) media. Clinical antibiotics were diluted in MHB2 media to 128 µg mL^−1^ and then diluted 2-fold in MHB2 media in a 96-well plate. 50 µL standardized cultures were added to each well containing 50 µL diluted drugs. The 96-well plate was sealed using Parafilm and placed in 37°C static incubator for 24 hours. 96-well plate was read at OD_600_ after 24 hours.

#### Time-kill assay

Overnight cultures were grown in lysogeny broth (LB) media at 37°C in shaking incubator (220 rpm) for 24 hours and synchronized 1:100 in LB media at 37°C in shaking incubator (220 rpm) for 3 hours and standardized to OD_600_ 0.01 in MHB1 media. 2-fold dilutions of bac7 peptide were performed in MHB1 media to achieve 0.5×, 1×, and 4× MIC concentrations, respectively. At each timepoint, 100 µL from culture tubes was transferred to 96-well plate and serially diluted 10-fold in 1× PBS. 5 µL serial dilutions were spot plated onto LB agar plate and placed in 37°C static incubator for 24 hours. Bacterial growth was assessed after 24 hours by enumeration of CFU mL^−1^. All time-kill assays were performed in biological triplicate with error reported as ± SEM.

### Biofilm viability assay

Clinical antibiotic minimal biofilm eradication concentration assays were performed by growing overnight cultures in lysogeny broth (LB) media at 37°C in shaking incubator (220 rpm) for 24 hours. Overnight cultures were standardized to OD_600_ 0.5 (9.75 × 10^9^) in biofilm media and 200 µL were pipetted into Calgary Biofilm Device (CBD) (Innovotech) 96-well plate with peg lid and placed in 37°C static incubator for 24 hours. Tigecycline, chloramphenicol, ertapenem, and gentamicin were diluted from 64 µg mL^−1^ to 0.5 µg mL^−1^ using 2-fold dilutions in new 96-well plate using Mueller Hinton Broth 2 (MHB2) media. Peg lid from CBD device was transferred to 96-well drug dilution plate, sealed with parafilm, and placed in 37°C static incubator for 24 hours. Following incubation with the antibiotics, the peg lid was placed into 96-well plate containing 200 µL 0.1% Crystal Violet (CV) for 15 minutes. The peg lid was removed and dried in fume hood for 24 hours. De-staining was completed by solubilizing the CV stain on the peg lid in 30% acetic acid for 15 minutes and the solubilized stain was read at OD_550_ using a BioTek SYNERGY H1 plate reader.

### Confocal microscopy z-stack imaging

#### Biofilm growth and treatment

Overnight cultures were grown in lysogeny broth (LB) media in shaking incubator (220 rpm) at 37°C for 24 hours. Overnight cultures were standardized to OD_600_ 0.5 in biofilm media and pipetted into #1.5 14 mm glass diameter Matsunami dishes (VWR). Matsunami dishes were sealed with parafilm and placed in static incubator at 37°C for 24 hours. After 24 hours, supernatant was removed from the biofilms and replaced with MHB1 media with 15 µmol L^−1^ bac7 or water only for no treatment control samples. Treatments performed for FITC-bac7 were performed as described above.

#### Staining

Post-24-hour treatment, supernatant of biofilms was removed, samples were washed with 1 mL phosphate buffer saline (PBS) and stained for imaging. To visualize the bacterial cells, the biofilms were stained with 5 µmol L^−1^ SYTO 9 green fluorescent stain in 1× PBS using a rocker at medium speed for 1 hour. SYTO 9 was removed, samples were washed with 1× PBS, and 50 µg mL^−1^ calcofluor white dye in molecular-grade water was added and rocked at medium speed for 5 minutes to stain the polysaccharide matrix. Calcofluor white dye was removed, and samples were washed with 1× PBS before confocal imaging. When visualizing biofilms treated with FITC-bac7, post-staining with calcofluor white stain was performed as described above. BactoView™ (red) cell stain was used in place of SYTO 9 at 5 µmol L^−1^ in 1× PBS with the same staining procedure as described above for SYTO 9.

#### Imaging

Biofilms were imaged using confocal z-stack imaging on a Zeiss LSM 710 confocal microscope with a 63× oil immersion lens. Lens oil was applied to the microscope lens, and sample dish (Matsunami Glass dish) was placed onto the microscope lens holder. Laser channels 488 and 543 nm laser channels for imaging of calcofluor white and SYTO 9 staining, respectively. When using BactoView™ laser channels 588 and 543 nm were used to visualize the red cell fluorescence and calcofluor white polysaccharide stain, respectively. Images were taken using confocal z-stacks (0-60 µm) and 3D renderings of the z-stack images were generated using ZEN microscope software.

#### BiofilmQ analysis

Confocal z-stack imaging files were processed with BiofilmQ software package using MATLAB_R2024b. Biofilm segmentation was done using a cube size of 1.32 µm resulting in 25 bins for MRSN 564304 and 15 bins for MRSN 1912. Fluorescence intensity as the bins increase in distance from the surface (dz (µm)) is measured to assess mean fluorescence through the layers of the biofilm. 4D XYZC plots were generated separately for red fluorescence and green fluorescence with the intensity of the respective signals displayed as heat map with the legend for each shown to the right.

### Skin abscess mouse model

Bacterial cultures were grown in tryptic soy broth (TSB) media until OD 0.5 was achieved. Cells were washed twice with 1× PBS and resuspended in 1× PBS to reach a final concentration of 1×10^4^ CFU mL^−1^. Six-week-old female CD-1 mice were anesthetized with isoflurane and subjected to a 1-cm-long superficial linear skin abrasion on their backs. 20 µL bacterial load resuspended in 1× PBS was inoculated over the abraded area. Polymyxin B control and bac7 were diluted in water to 10× the minimum inhibitory concentration (MIC) and applied to the infected region 5 hours post-infection. The animals were euthanized 2 and 4 days after infection, skin samples and organs were collected and homogenized to enumerate the bacterial burden. Scarified skin areas and organ systems were excised and homogenized using a bead beater (25 Hz) for 20 minutes and serially diluted using 10-fold dilutions in 1× PBS for CFU quantification. Six mice per group (n = 6) were used for the experimental groups and statistical significance was determined using one-way ANOVA, p-values are shown for each of the groups, with all groups compared to the untreated control group. Mice were single-housed to avoid cross-contamination and maintained under a 12-hour light/dark cycle at 22 °C with controlled humidity at 50%. The skin abscess infection mouse model was revised and approved by the University Laboratory Animal Resources (ULAR) from the University of Pennsylvania (Protocol 806763).

## Acknowledgments

This work was supported by National Institute of Allergy and Infectious Diseases of the National Institutes of Health under award number R00AI163295 awarded to R.M.F. C.F.-N. holds a Presidential Professorship at the University of Pennsylvania and acknowledges funding from the Procter & Gamble Company, United Therapeutics, a BBRF Young Investigator Grant, the Nemirovsky Prize, Penn Health-Tech Accelerator Award, Defense Threat Reduction Agency grants HDTRA11810041 and HDTRA1-23-1-0001, and the Dean’s Innovation Fund from the Perelman School of Medicine at the University of Pennsylvania. Research reported in this publication was supported by the Langer Prize (AIChE Foundation), the NIH R35GM138201, and DTRA HDTRA1-21-1-0014.

## Author Contributions

The transcriptional RNA sequencing and qRT-PCR analysis was performed by BVM. The confocal microscopy biofilm experiments and time-kill assays were performed by RLB. The membrane potential assays were performed by RLB and FZS. The MIC and MBEC_90_ experiments were performed by RLB and GNE. The fluorescence microscopy was performed by LS and PE. The murine skin abscess model was performed by MDTT and CFN. The fimbriae analysis was performed by SG, PLW, and DAR. RMF and RLB wrote the manuscript with input from others.

